# Placental DNA methylation levels at *CYP2E1* and *IRS2* are associated with child outcome in a prospective autism study

**DOI:** 10.1101/501007

**Authors:** Yihui Zhu, Charles E. Mordaunt, Dag H. Yasui, Ria Marathe, Rochelle L. Coulson, Keith W. Dunaway, Cheryl K. Walker, Sally Ozonoff, Irva Hertz-Picciotto, Rebecca J. Schmidt, Janine M. LaSalle

## Abstract

DNA methylation acts at the interface of genetic and environmental factors relevant for autism spectrum disorder (ASD). Placenta, normally discarded at birth, is a potentially rich source of DNA methylation patterns predictive of ASD in the child. Here, we performed whole methylome analyses of placentas from a prospective study of high-risk pregnancies. 400 differentially methylated regions (DMRs) discriminated placentas stored from children later diagnosed with ASD compared to typical controls. These ASD DMRs were significantly enriched at promoters, mapped to 596 genes functionally enriched in neuronal development, and overlapped genetic ASD risk. ASD DMRs at *CYP2E1* and *IRS2* reached genome-wide significance, replicated by pyrosequencing, and correlated with expression. Methylation at *CYP2E1* associated with both ASD diagnosis and *cis* genotype, while methylation at *IRS2* was unaffected by *cis* genotype but modified by preconceptional maternal prenatal vitamin use. This study therefore identified two potentially useful early epigenetic markers for ASD in placenta.

## Introduction

Autism spectrum disorder (ASD) is a heterogeneous neurodevelopmental disorder diagnosed by a combination of behavioral features including restricted interests, repetitive behaviors, language deficits, and impairments in social communication (Baio et al., 2018). 1 in 59 children in the United States are diagnosed with ASD, at a mean age of 4.2 years (Baio et al., 2018). ASD is currently diagnosed by clinicians trained on the Autism Diagnostic Observation Schedule (ADOS) and the Autism Diagnostic Interview - Revised (ADI-R) according to the Statistical Manual of Mental Disorders (DSM-5) which is most accurate at or after 36 months (Baio et al., 2018). However, an early assessment of ASD risk could identify infants and toddlers who would benefit from behavioral interventions that improve cognitive, social, and language skills.

Monozygotic versus dizygotic twin and sibling studies suggest a strong genetic basis for ASD (Hannon et al., 2018; M. B. Jones & Szatmari, 1988; Ritvo et al., 1989; Tsai & Bell, 2015; Wessels & Pompe van Meerdervoort, 1979). However, mutations in any individual gene account for less than 1% of ASD cases (Bourgeron, 2015; Tsai & Bell, 2015). Genetic sequencing analyses can only identify a potentially causative genetic abnormality in ~25% of clinical ASD diagnoses (Bourgeron, 2015; Tsai & Bell, 2015; Turner et al., 2016). While genome-wide association studies (GWAS) also support common genetic variants in ASD, the complexity and heterogeneity of ASD has been a major challenge (Abrahams et al., 2013; Grove et al., 2017; Iossifov et al., 2014; Sanders et al., 2015). Evidence for environmental risk factors in ASD point to *in utero* maternal exposures such as air pollution, fever, or asthma and nutrients specifically the absence of pre-conceptional prenatal vitamin intake (Raz et al., 2015; Schmidt et al., 2011, 2012; Zerbo et al., 2013). Maternal prenatal vitamins, which contain high levels of folate and other additional B vitamins, protect offspring by up to 70% for neural tube defects (Caramaschi et al., 2017; Howsmon, Kruger, Melnyk, James, & Hahn, 2017; Kalkbrenner, Schmidt, & Penlesky, 2014; Relton et al., 2004; Rush, Katre, & Yajnik, 2014; Zeisel, 2009), and correlate with an overall 40% reduction in ASD risk if taken during the first month of pregnancy (P1) (Schmidt et al., 2011, 2012; Suren et al., 2013). This finding was replicated with a large prospective study in Norway including over 80,000 pregnancies (Suren et al., 2013).

DNA methylation shows dynamic changes during fetal development (Vogel Ciernia & LaSalle, 2016; Crawley, Heyer, & LaSalle, 2016; Smallwood & Kelsey, 2012b) and contains the molecular memory of *in utero* experiences such as maternal nutrition (Howsmon et al., 2017; Teh et al., 2014). Term placenta is an accessible fetal tissue that maintains the distinctive embryonic bimodal DNA methylation pattern, in which expressed genes are marked by higher methylation levels (Schmidt et al., 2016; Schroeder et al., 2013, 2015, 2016). Placenta therefore offers a unique window to study DNA methylation patterns that may reflect altered fetal development relevant to ASD genetic risk (Schroeder et al., 2015; Smallwood & Kelsey, 2012b, 2012a; Watson & Cross, 2005). Specifically, a recent study of polygenic risk scores for schizophrenia demonstrated a significant interaction of genetic risk with maternal perinatal environmental factors that affected placental gene expression (Ursini et al., 2018). Previous analyses of DNA methylation patterns in placenta samples from a high-risk ASD cohort also identified an association between self-reported use of lawn and garden pesticides and large-scale changes in DNA methylation patterns, and identified a putative enhancer of the *DLL1* gene as differentially methylated in ASD (Schmidt et al., 2016; Schroeder et al., 2016).

Here, we continue the epigenetic investigation of ASD risk through the novel approach of identifying differentially methylated regions (DMRs) in whole methylomes from placenta samples from male children later diagnosed with ASD compared to children with typical development (TD) controls. Two genome-wide significant ASD-associated DMRs at *CYP2E1* and *IRS2* were further validated and investigated for effects of genotype, RNA expression, and protein levels as well as interactions with preconception prenatal vitamin use. Understanding the epigenetic patterns of ASD associated with maternal prenatal vitamin use in placenta could lead to the development of preventative and therapeutic early interventions for high-risk children with ASD.

## Results

### Placenta ASD DMRs discriminate ASD from TD samples

To identify novel differentially methylated gene loci between ASD and TD, a differentially methylated region (DMR) bioinformatic analysis was performed on placenta whole genome bisulfite sequencing (WGBS) data (Schmidt et al., 2016; Schroeder et al., 2016). 400 DMRs were identified with a threshold of > 10% methylation difference between ASD and TD groups, and these were associated with 596 genes (**Fig. 1A**, **Supplementary Table 1**). There was no bias for gene length in the ASD DMR associated genes compared to all human genes (**Supplementary Fig. 1**). 296 DMRs were hypermethylated, while 104 DMRs were hypomethylated in ASD compared to TD placenta (**Fig. 1A**). Principal component analysis (PCA) using methylation levels for each sample over the 400 DMRs demonstrated a clear separation of placental samples by child outcome of ASD versus TD (**Fig. 1B**). In addition, most ASD DMRs showed a highly significant association with Mullen Scales of Early Learning (MSEL) scores and autism severity score from Autism Diagnostic Observation Schedule (ADOS), but not with potential confounding variables (**Supplementary Table 2**, **Supplementary Fig. 2**).

**Figure 1.**
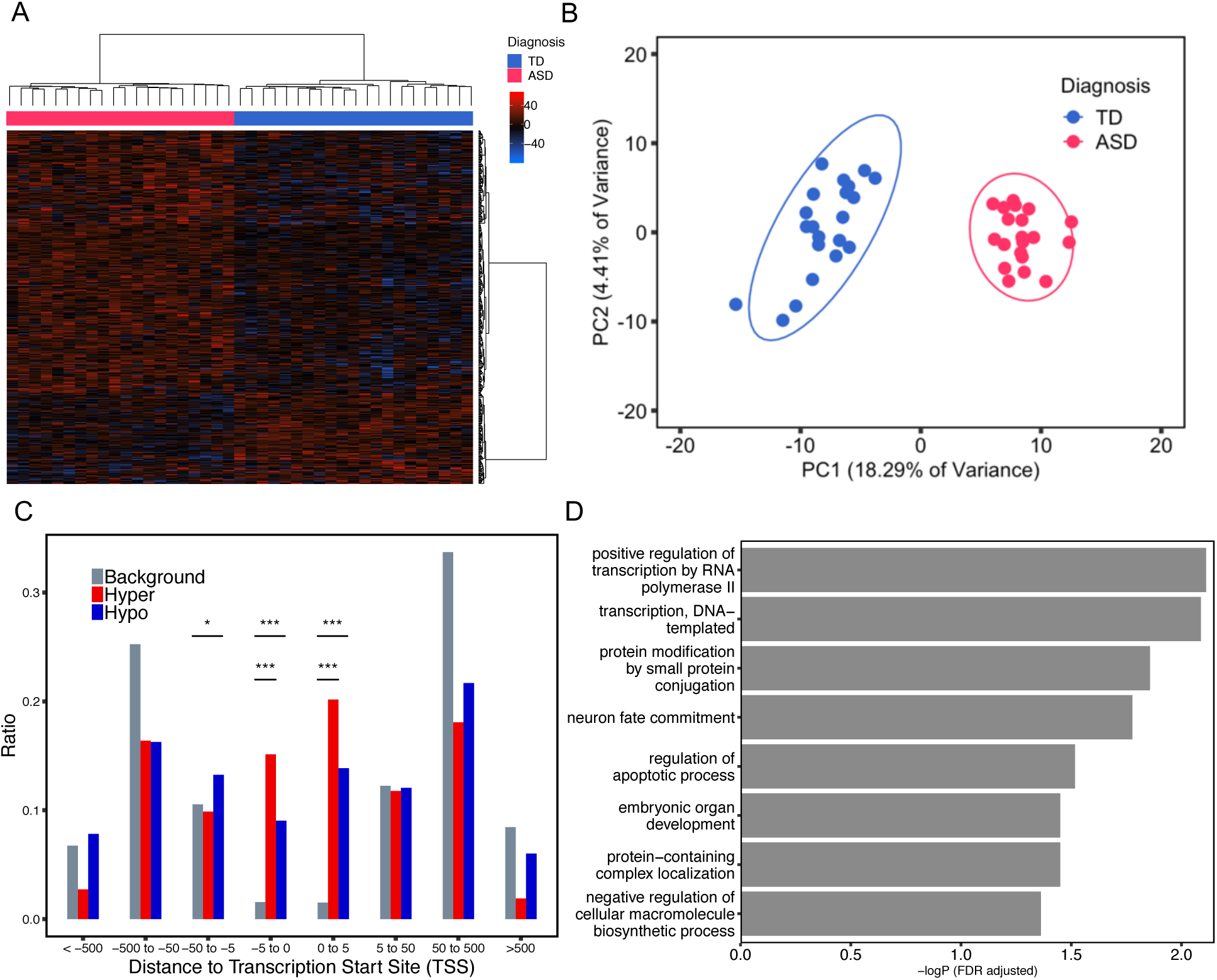
Differentially methylated regions (DMRs) in placenta distinguished ASD from typical development (TD) child outcomes. **A**. Heatmap and hierarchical clustering of 20 ASD versus 21 TD placental samples using methylation levels in the 400 identified ASD DMRs. Percent methylation for each sample relative to the mean methylation at each ASD DMR was plotted as a heatmap, with black representing no difference, hyper-methylation (red) and hypo-methylation (blue). Columns were clustered by child outcome, ASD (red) or TD (blue), while rows were clustered by methylation direction. **B**. Principal component analysis (PCA) of TD vs ASD placental samples on the basis of methylation at 400 ASD DMRs. Ellipses represent the 95% confidence interval for each group. The non-overlapping ellipses showed a significant difference between ASD and TD for these DMRs’ methylation level (*p* < 0.05). **C**. Location relative to genes for hypermethylated (red) or hypomethylated (blue) ASD DMRs compared to background (grey). Distributions of locations relative to transcription start sites (TSS) are shown on the x-axis. The ratio plotted on the y-axis is calculated by the number of genes at each binned location divided by the total number of genes (**Supplementary Table 2**). \**p* < 0.05, \*\**p* < 0.01, \*\*\**p* < 0.001 by Fisher’s exact test. **D**. Bar graph represents the significant results from gene ontology and pathway enrichment analysis of ASD DMRs associated genes compared to background by Fisher’s exact test (FDR adjusted -log *p*-value, x-axis).

### Placenta ASD DMRs were enriched for transcription start sites and genes that function in transcriptional regulation and neuronal fate

To further study the location and function of ASD DMRs in placenta, we calculated the location of each ASD DMR relative to the assigned gene’s transcription start site (TSS) (**Supplementary Table 3**). Both hyper- and hypomethylated ASD DMRs were enriched within 5kb on either side of TSS compared with background regions (**Fig. 1C**). Gene ontology (GO) analysis of ASD DMRs genes revealed significant enrichment for functions in transcription, protein modification, embryonic organ development, and neuron fate commitment by Fisher’s exact test after false discovery rate (FDR) multiple test correction (**Fig. 1D, Supplementary Table 4**).

### Placenta DMR genes were enriched in ASD but not ID risk genes

To test a hypothesized overlap between epigenetic and genetic ASD risk loci observed previously in ASD and neurodevelopmental disorder brain tissues (Vogel Ciernia et al., 2018), we investigated the possible overlap of placenta ASD DMR genes with identified genetic risk factors for ASD and other types of intellectual disability (ID). First, the curated Simons Foundation Autism Research Initiative (SFARI) gene list was separated into six categories based on SFARI ASD gene scores (Abrahams et al., 2013). The entire list of SFARI genes as well as the high confidence gene list both showed significant overlap with placenta ASD DMR genes (**Fig. 2A**, **Supplementary Table 5**). The 39 genes in common between the SFARI gene list and placenta ASD DMRs were significantly enriched for functions in positive regulation of histone H3K4 methylation, multicellular organ development, and system development. Second, high risk ASD genes from Sanders *et al*. (Sanders et al., 2015) and likely gene-disrupting (LGD) recurrent mutations and missense mutation on *de novo* mutations to ASD gene lists from whole genome exome sequencing (Iossifov et al., 2014) were also significantly enriched for placental ASD DMRs. In contrast, no significant enrichment was observed for placental ASD DMRs with ID, Alzheimer’s disease, or lung cancer genetic risk (Gilissen et al., 2014) or a random set of 400 genomic regions mapped to 600 genes (**Fig. 2A**, **Supplementary Table 5**). When placental ASD DMRs were separated by direction of change, hypomethylated ASD DMRs exhibited more categories of significant enrichment with ASD genetic risk compared with hypermethylated ASD DMRs (**Supplementary Fig. 3**)

**Figure 2.**
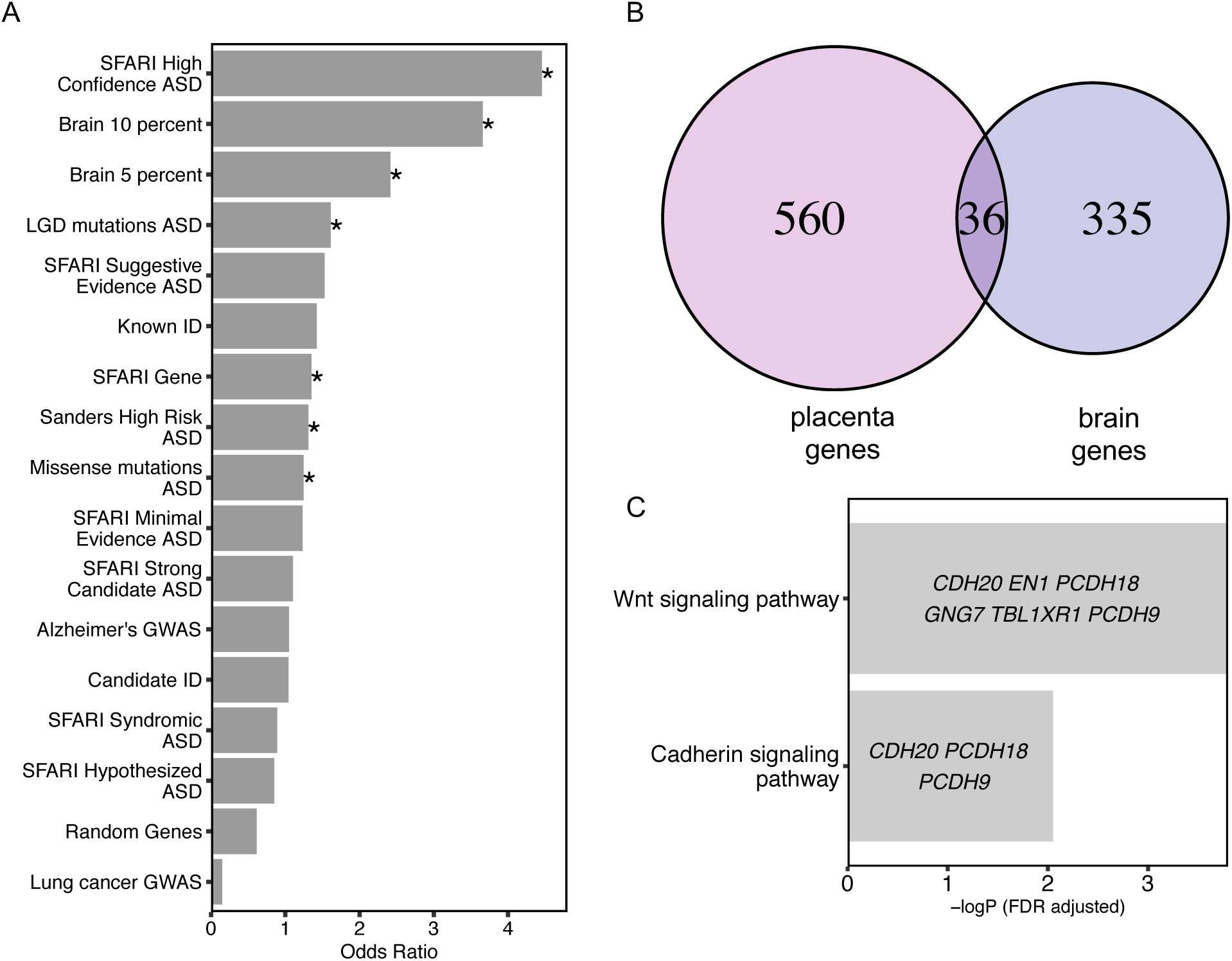
Placenta ASD DMR genes overlapped with ASD DMR associated genes from postmortem brain and known genetic risk for ASD, but not for other disorders. **A**. Placenta ASD DMR associated genes were compared for significant overlap with ASD DMR genes identified from ASD postmortem brain (Vogel Ciernia et al, 2018, based on 10% or 5% methylation difference cutoffs), as well as multiple curated gene lists of ASD, intellectual disability, or unrelated disorder genetic risks, or a randomly generated gene list (\**p* < 0.05 FDR corrected two-tailed Fisher’s exact test, ranked by odds ratio). SFARI: Simons Foundation Autism Research Initiative (Abrahams et al., 2013), LGD: likely gene disrupting mutation, ASD: autism spectrum disorder, Alzheimer: Alzheimer’s Disease, ID: intellectual disability. **B**. Venn diagram represents the overlap of 36 genes associated with placenta ASD DMRs and brain ASD DMRs based on 10% methylation differences between ASD versus TD (**Supplementary Table 5**).

Methylation data from human postmortem brain was obtained from previous published datasets, GSE8154 (ASD and TD) (Vogel Ciernia et al., 2018). **C**. Gene ontology and pathway analysis on the 36 genes in common between placenta ASD DMRs and brain ASD DMRs associated genes. Enrichment tests were done on Fisher’s exact test with FDR 0.05 correction. Genes in each gene ontology term are shown within each bar.

### Placenta ASD DMR genes significantly overlapped with brain ASD DMRs that were enriched for Wnt and cadherin signaling pathways

From a prior methylation analysis in brain frontal cortex, 210 ASD discriminating DMRs from brain (10% methylation difference between ASD and TD) were identified, which mapped to 371 genes (Vogel Ciernia et al., 2018). A significant overlap of ASD DMR genes was observed between placenta and brain (Fisher’s exact test, *p*-value < 0.001), with 36 genes in commons (**Fig. 2B**, **Supplementary Table 5**). Those 36 genes were significantly enriched for functions in the Wnt signaling and cadherin signaling pathways (**Fig. 2C**). Of these shared genes four overlapped with SFARI genetic risk: *GADD45B, MC4R, PCDH9*, and *TBL1XR1*.

### Placenta ASD DMRs were enriched for placental and brain active promoter H3K4me3 peaks, promoter flanking regions, and CpG shores

To functionally annotate the ASD DMRs identified by placenta WGBS, multiple histone modification ChIP-seq peaks and chromatin state predictions from multiple tissue types in the Roadmap Epigenomics Projects were compared to ASD DMR chromosomal locations for enrichment compared to background regions (Kundaje et al., 2015). Placental ASD DMRs were significantly enriched for H3K4me3 and H3K4me1 marks of promoters and enhancers across multiple tissues, although placental H3K4me3 marks showed the strongest (odds ratio = 17.08, FDR q < 1.8E-42) and brain H3K4me3 marks showed the second strongest enrichment (odds ratio = 13.75, FDR q < 3.55E-31) (**Fig. 3A**). Next, we overlapped ASD DMRs with published chromatin state predictions that use histone modification ChIP-seq data to annotate the genome into 15 functional states (chromHMM) (Ernst & Kellis, 2012). Placental ASD DMRs showed significant enrichment in regions flanking transcription start site (TssAFlnk) and transcription start site (TssA) compared to background over multiple tissues (**Fig. 3B**). When separated by directional change in ASD, both hyper- and hypomethylated ASD DMRs were significantly enriched for H3K4me3 peaks, transcription start sites and their flanking regions, as well as enhancers (**Supplementary Fig. 4**).

**Figure 3.**
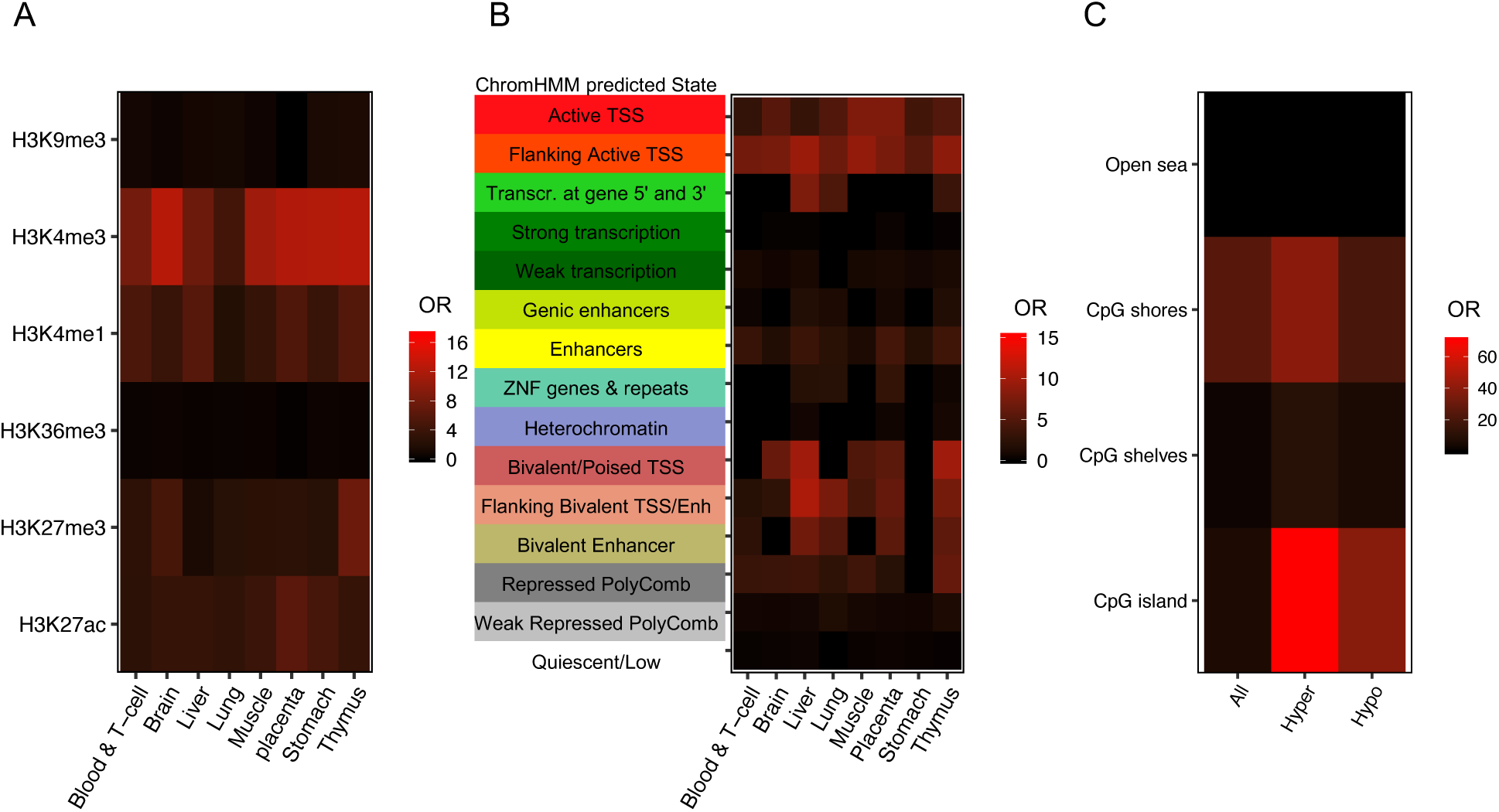
Placenta ASD DMRs were enriched at H3K4me3 regions, flanking promoter regions, and CpG shores. **A**. Placenta ASD DMRs were examined for enrichment with histone modification ChIP-seq peaks from the Epigenome Roadmap using the LOLA package. Enrichments are plotted as the odds ratio) in a heat map for each of 8 different tissue types and 6 types of modified histone marks (Sheffield & Bock, 2016). **B**. Enrichment tests on chromatin states from chromHMM categories in the Epigenome Roadmap and placental ASD DMRs from this study were performed using LOLA, with each row representing a different ChromHMM predicted state and each column a single tissue type. **C**. Placenta ASD DMRs (categorized as all, hypermethylated, or hypomethylated in ASD) were tested for enrichment based on CpG island location. The human genome was separated into CpG islands, CpG shores, CpG shelves and open sea.

We also separated the genome into four parts relative to CpG island location (Aryee et al., 2014; Sandoval et al., 2011; Timp et al., 2014). CpG shores were defined as the region within 2 kb on both sides of CpG islands, while another 2 kb extension from the shores were defined as CpG shelves. The remaining regions were defined as “open sea”. Placental ASD DMRs showed significant enrichment at CpG shores, and hypermethylated ASD DMRs more significantly overlapped CpG islands compared with hypomethylated DMRs (**Fig. 3C**).

### Two genome-wide significant placental ASD DMRs at *CYP2E1* and *IRS2* validate by pyrosequencing and correlated with gene expression

Two of the 400 ASD DMRs identified in ASD placenta reached genome-wide significance by family-wide error rate (FWER), including chr10: 133527713–133529507, located inside *CYP2E1* (cytochrome P450 2E1), and chr13: 109781111–109782389 located inside *IRS2* (insulin receptor substrate 2) (**Fig. 4**). The *CYP2E1* DMR was located after the first exon, included the first intron and part of the second exon, and was hypomethylated in ASD versus TD (**Fig. 4A**). The *IRS2* DMR, spanning the end of the first exon and the beginning of first intron and was hypermethylated in ASD versus TD (**Fig. 4B**). Both *CYP2E1* and *IRS2* were also present in the gene lists overlapping with brain ASD DMR related genes and high risk ASD genes (**Fig. 2A, Supplementary Table 5**) (Sanders et al., 2015).

**Figure 4.**
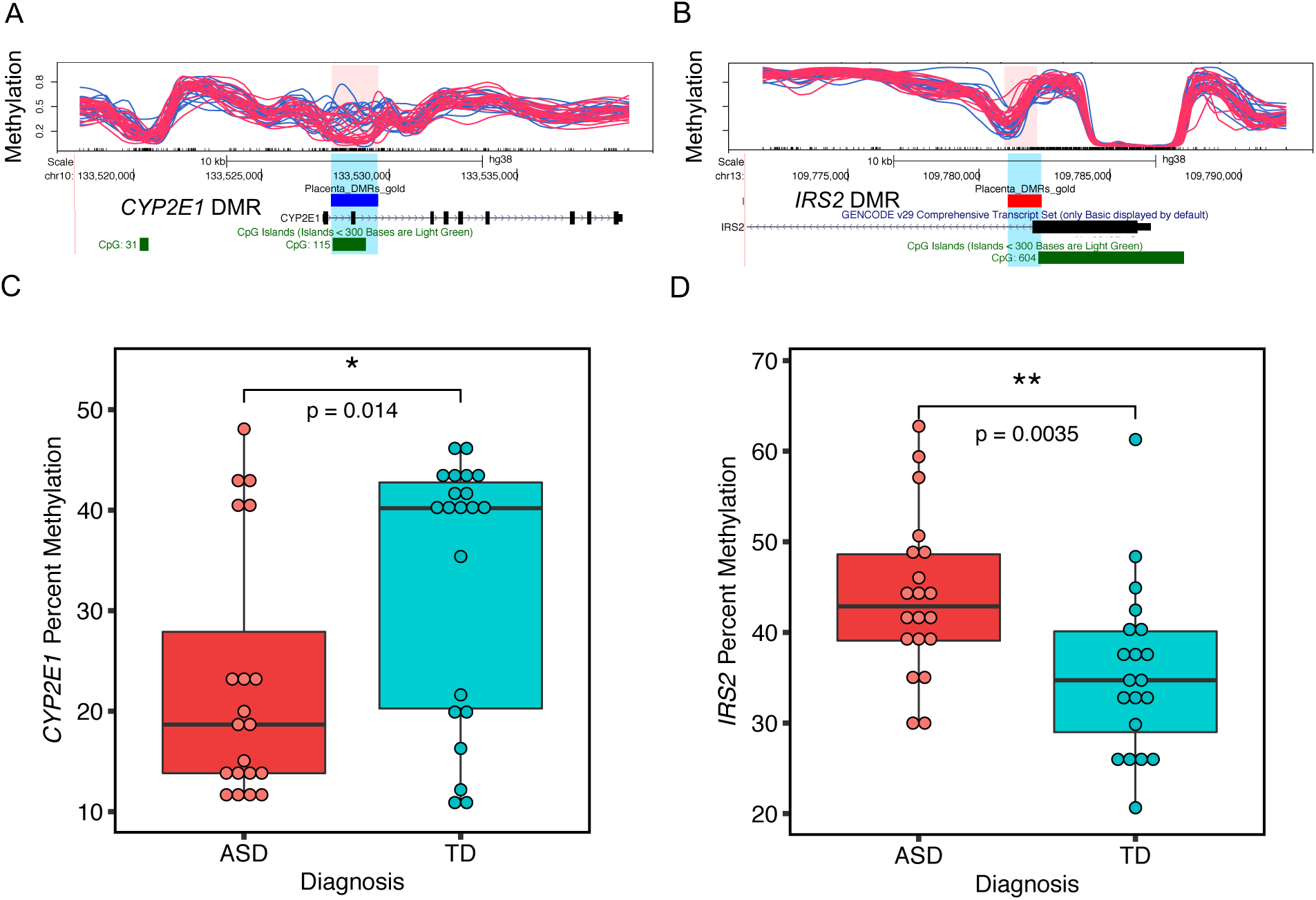
Two genome-wide significant placental DMRs located at *CYP2E1* and *IRS2* were validated by pyrosequencing. **A** and **B** show the location relative to genes, and CpG islands of the two genome-wide significant DMRs (highlighted in pink and blue) in the UCSC Genome Browser. In the upper tracks, each line represents percent methylation (y-axis) of a single individual by WGBS analysis. Blue lines represent TD and red lines represent ASD samples. **A**. Hypomethylated DMR at *CYP2E1* with 10 kb upstream and 10 kb downstream. **B**. Hypermethylated DMR at *IRS2* with 10 kb upstream and 10 kb downstream. **C**. The *CYP2E1* DMR percent methylation was significantly associated with child outcome and verified by pyrosequencing (two-tailed t-test, *p*-value = 0.014). The y-axis represents the average percent DNA methylation across the DMR regions from pyrosequencing. Each dot represented one sample. **D**. Pyrosequencing validation on *IRS2* DMR’s methylation with child outcome (two-tailed t-test, *p*-value = 0.0035). \**p* < 0.05, **p < 0.01

Pyrosequencing was performed as an independent method to verify methylation differences between ASD and TD placental samples at *CYP2E1* and *IRS2* DMRs (**Supplementary Table 6**). For the *CYP2E1* DMR, there was a significant difference in average percent methylation detected by pyrosequencing between ASD and TD samples (**Fig. 4C**). 13 CpG sites were included in the *CYP2E1* DMR pyrosequencing test, and all but two also showed individually significant differences between ASD and TD after false discovery rate (FDR) correction (**Supplementary Table 6**, **Supplementary Fig. 5A**). Pyrosequencing also confirmed a significant difference between ASD and TD average percent methylation at the *IRS2* DMR (**Fig. 4D**) and all of the 11 CpG sites individually assayed at *IRS2* (**Supplementary Table 6**, **Supplementary Fig. 5B**).

While MARBLES placenta samples were not collected in a manner conducive to RNA stability for gene expression analyses, we were able to examine expression level of both *CYP2E1* and *IRS2* in MARBLES umbilical cord blood from an Affymetrix Human Gene 2.0 array analysis in a related study (Mordaunt, Park, et al., 2018). A trend for lower *CYP2E1* transcript levels was observed in ASD compared with TD cord blood samples (**Fig. 5A**) consistent with the direction of the placental methylation for this locus (**Fig. 4C**). Similarly, a trend for higher *IRS2* expression in ASD versus TD cord blood was observed along with higher ASD methylation in placenta (**Fig. 4D**, **Fig. 5B**). To investigate the direction of expression changes for these genes in ASD brain, we utilized the dbMDEGA database (Zhang et al., n.d.). *CYP2E1* showed a significantly downregulated in ASD compared with TD in human male cortex, while *IRS2* trended for higher levels in ASD compared with TD. These were both in the same direction as ASD cord blood expression and ASD placental methylation compared to controls. Furthermore, a trend for higher IRS2 protein in ASD placenta samples compared with TD placenta samples was observed by Western blot (**Fig. 5C**, **Supplementary Fig. 6**), as expected based on transcript and methylation levels. Because of the distinctive methylation landscape in placenta, positive correlations between methylation and expression were expected for gene body locations outside of CpG islands (Schroeder et al., 2013).

**Figure 5.**
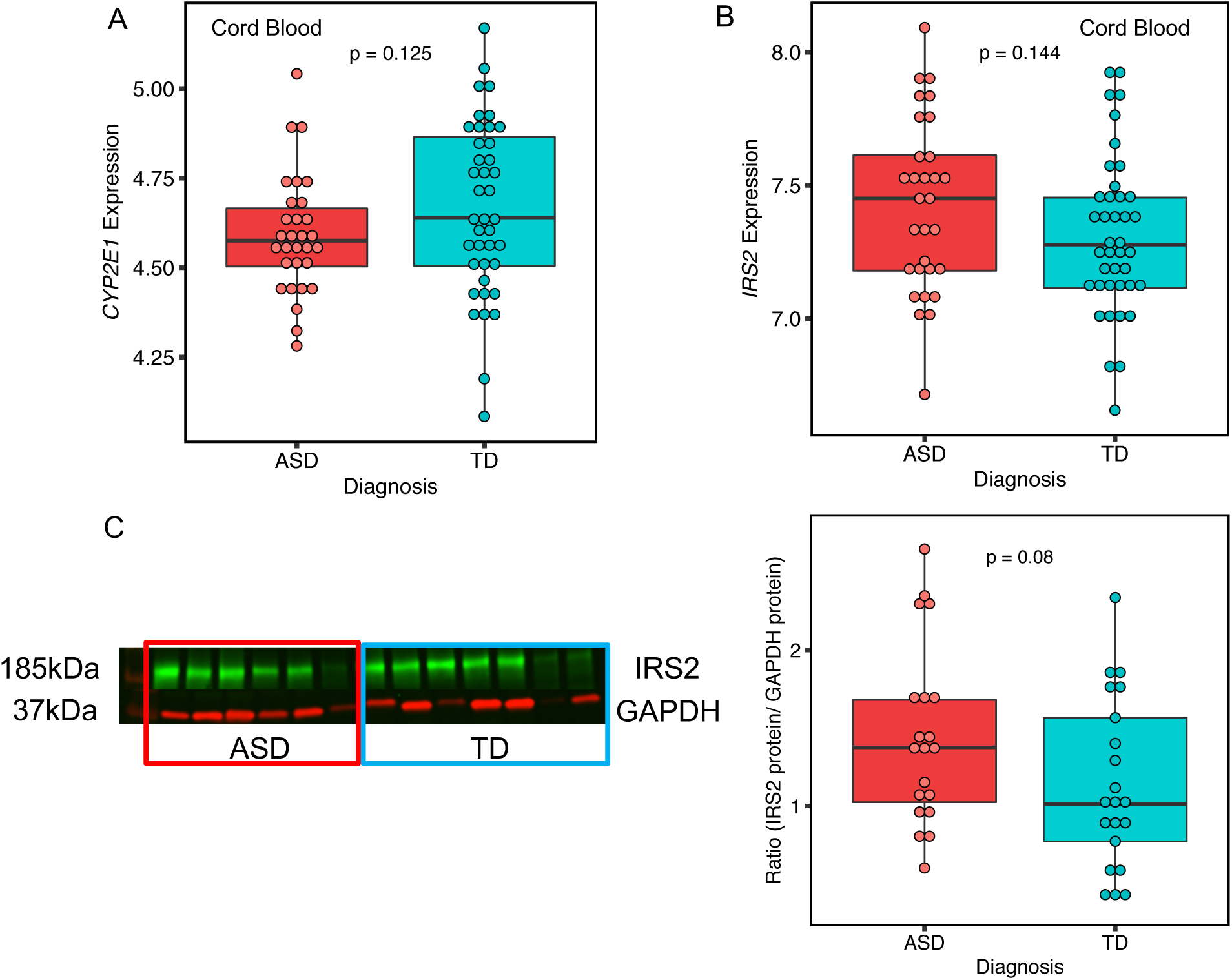
For both placental ASD DMRs at *CYP2E1* and *IRS2*, expression trended towards positive correlation with methylation. **A**. 30 ASD and 40 TD umbilical cord blood sample in MARBLES were included in this analysis. Affymetric array matrix data on the probe 16711001 was used to represent the expression of *CYP2E1* on the y-axis. Each dot was used to represent one individual (two-tailed t-test, *p*-value = 0.125). **B**. The same umbilical cord blood samples were used for measuring *IRS2* expression at the probe 16780917 (two-tailed t-test, *p*-value = 0.144). C. Representative Westerns blots are shown for the ratio of IRS2 to GAPDH (normalization control) in all 41 placenta samples of ASD and TD comparison with each dot representing one sample (two-tailed t-test, *p*-value = 0.08). A Western blot with 6 samples in ASD and 7 samples TD were showed at the left panel. IRS2 protein was labeled with green fluorescence at 185 kDa and GAPDH was marked with red fluorescence at 37 kDa.

### *CYP2E1* but not *IRS2* DMR methylation levels were associated with *cis* genotypes

We performed Sanger sequencing within the *CYP2E1* and *IRS2* ASD DMRs to identify single nucleotide polymorphisms (SNPs) that could explain some of the methylation differences. Two SNPs (rs943975, rs1536828) were identified within the boundaries of the *CYP2E1* DMR in the 41 placenta samples (**Supplementary Table 7**). A significant association between rs1536828 (but not rs943975) genotype and *CYP2E1* DMR percent methylation was observed, with samples homozygous for the minor allele (G/G) showing the lowest methylation (**Fig. 6A)**. A single informative SNP (rs9301411) was also identified within the *IRS2* DMR (**Supplementary Table 7**) but was not significantly associated with methylation level (**Fig. 6B**).

**Figure 6.**
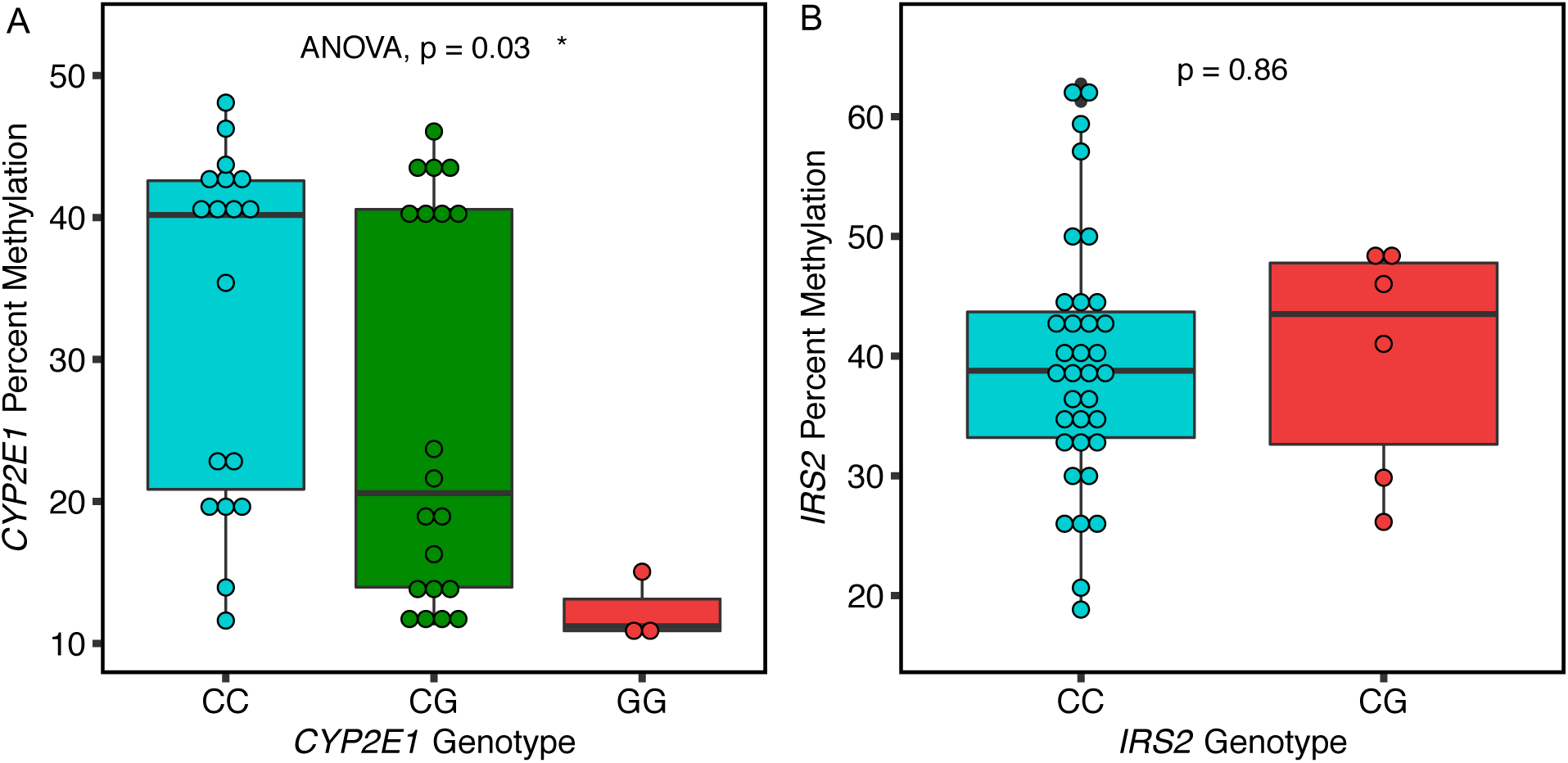
*Cis* genotype was significantly associated with *CYP2E1* but not *IRS2* DMR methylation levels. **A**. *CYP2E1* genotype at rs1536828 within the ASD DMR was significantly associated with *CYP2E1* DMR average percent methylation tested by ANOVA (*p*-value = 0.03). **B**. *IRS2* genotype at rs9301411 within the ASD DMR was not significantly associated with *IRS2* DMR methylation by two-tailed t-test (*p*-value = 0.86).

### Preconception prenatal vitamin use corresponded to protective placental DNA methylation patterns at *CYP2E1, IRS2*, and genome-wide

Placental samples from mothers who took prenatal vitamins during the first month of pregnancy showed a trend for higher *CYP2E1* DMR methylation that was not significant, but in the same direction expected for protection from ASD (**Fig. 7A**). At the *IRS2* DMR, however, there was a significant association with maternal prenatal vitamin use and lower methylation, also in ASD-protective direction (**Fig. 7B**).

To further explore the relationship between placental methylation patterns influenced by prenatal vitamin use in the first month of pregnancy, placental methylomes were analyzed for DMRs by prenatal vitamin use in the first month of pregnancy (PreVitM1) with more than 10% methylation difference, and 376 DMRs were identified over 587 genes (**Supplementary Table 8**). 60 genes overlapped between PreVitM1 DMRs and ASD DMRs in placenta (**Supplementary Table 8**, **Fig. 7C**). Gene ontology analysis showed that genes common to PreVitM1 and ASD DMRs were significantly enriched for functions in neuron fate commitment, transcription regulation, central nervous system development, and regulation of phosphatidylinositol 3-kinase activity (**Fig. 7D**). We also separated placental samples based on when mothers started taking prenatal vitamins during pregnancy into three periods (**Supplementary Table 9**). For the *CYP2E1* DMR, we found that during all three periods, ASD placental samples showed lower percent methylation on the *CYP2E1* DMR compared to TD (**Supplementary Fig. 7A**). The expected opposite finding of higher methylation levels in ASD compared with TD placental samples was observed at the *IRS2* DMR (**Supplementary Fig. 7B**).

**Figure 7.**
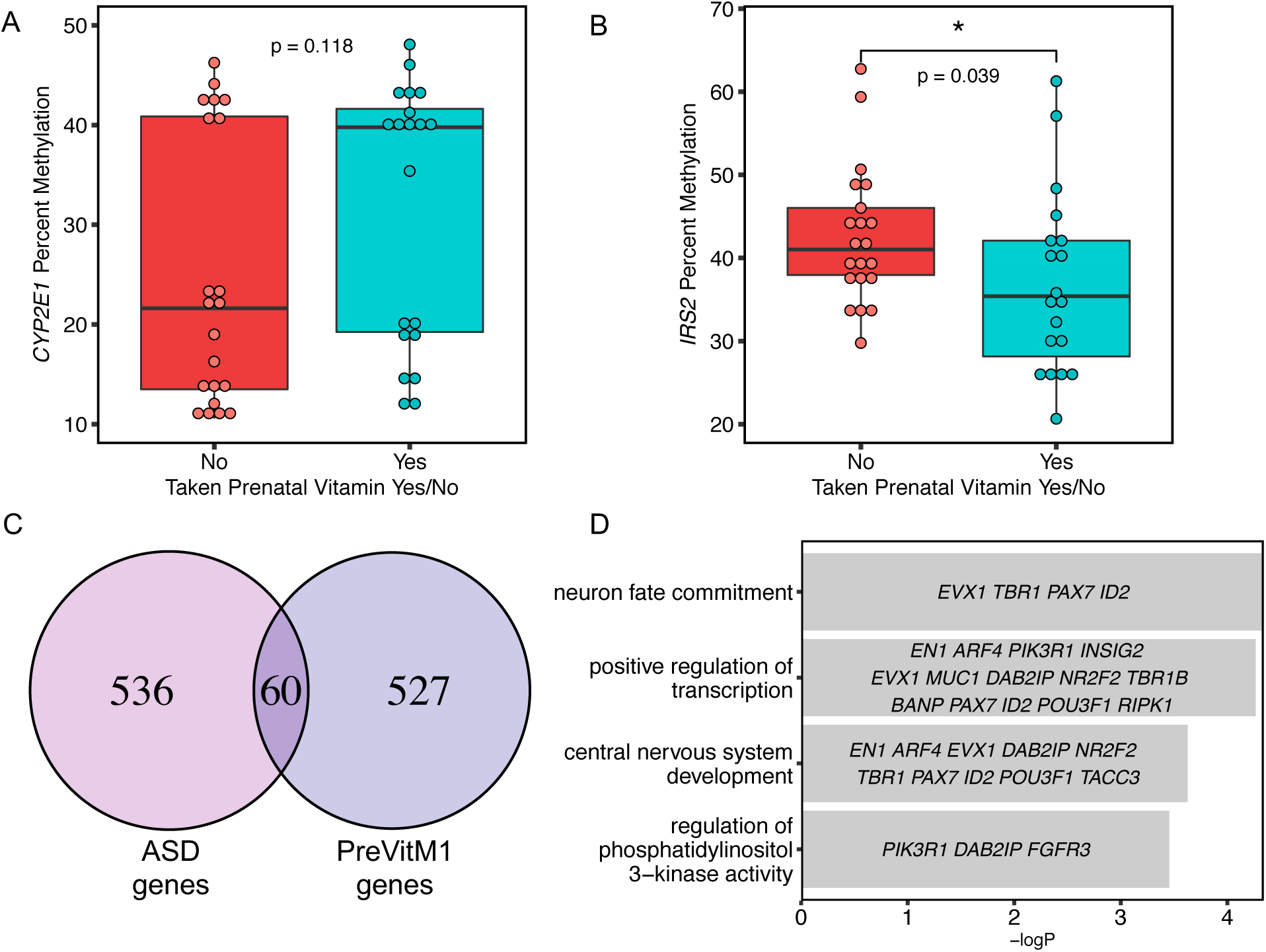
Preconception prenatal vitamin use was a significant modifier of *IRS2* methylation and associated DMRs overlapped ASD DMRs in placenta. For **A** and **B**, the x-axis represents maternal prenatal vitamins use during the first month pf pregnancy, while the y-axis shows the percent methylation. **A**. *CYP2E1* DMR methylation was not significantly altered by P1 prenatal vitamin use, although a trend was observed in the protective direction for ASD (two-tailed t-test, *p*-value = 0.118). **B**. Higher percent methylation at *IRS2* DMR was significantly associated with not taking prenatal vitamins at P1 (two-tailed t-test, *p*-value = 0.039), which is in the same direction as ASD risk. **C**. DMRs identified based on P1 prenatal vitamins use were associated with 587 genes, which showed a significant overlap with ASD DMR associated genes (Fisher’s exact test, *p*-value < 2.528e-16). **D**. Gene ontology and pathway analysis was performed on the overlapped gene list (60 genes) (**Supplementary Table 8**) between placenta ASD DMR and P1 prenatal vitamin DMR associated genes for enrichment by Fisher’s exact test with -log (*p*-value) represented on the x-axis. Genes in each gene ontology (GO) term are shown within each bar.

To further investigate the potential inter-relatedness of diagnosis, prenatal vitamin use, and *cis* genotype on methylation at *CYP2E1* and *IRS2* DMRs, we calculated associations between each factor and methylation separately by two-tailed t-test or ANOVA, as well as two-way diagnosis and PreVitM1; diagnosis and genotype; genotype and PreVitM1 by Pearson’s chi-squared test. These analyses illustrate that *CYP2E1* genotype and diagnosis significantly contributed to *CYP2E1* DMR methylation, while PreVitM1 and diagnosis were significantly associated with *IRS2* DMR methylation (**Fig. 8A, Fig. 8B**). No significant association was found between two-way interactions among each of the three factors and each DMR methylation level by ANOVA (**Supplementary Table 10**). When combining *CYP2E1* genotype, PreVitM1, and diagnosis to predict methylation at the *CYP2E1* DMR, a significant association was observed on *CYP2E1* DMR methylation with diagnosis and rs1536828 genotype (**Fig. 8C**, **Supplementary Fig. 8A**). At the *IRS2* DMR, PreVitM1 and diagnosis significantly contributed to *IRS2* DMR methylation (**Fig. 8D**, **Supplementary Fig. 8B**).

**Figure 8.**
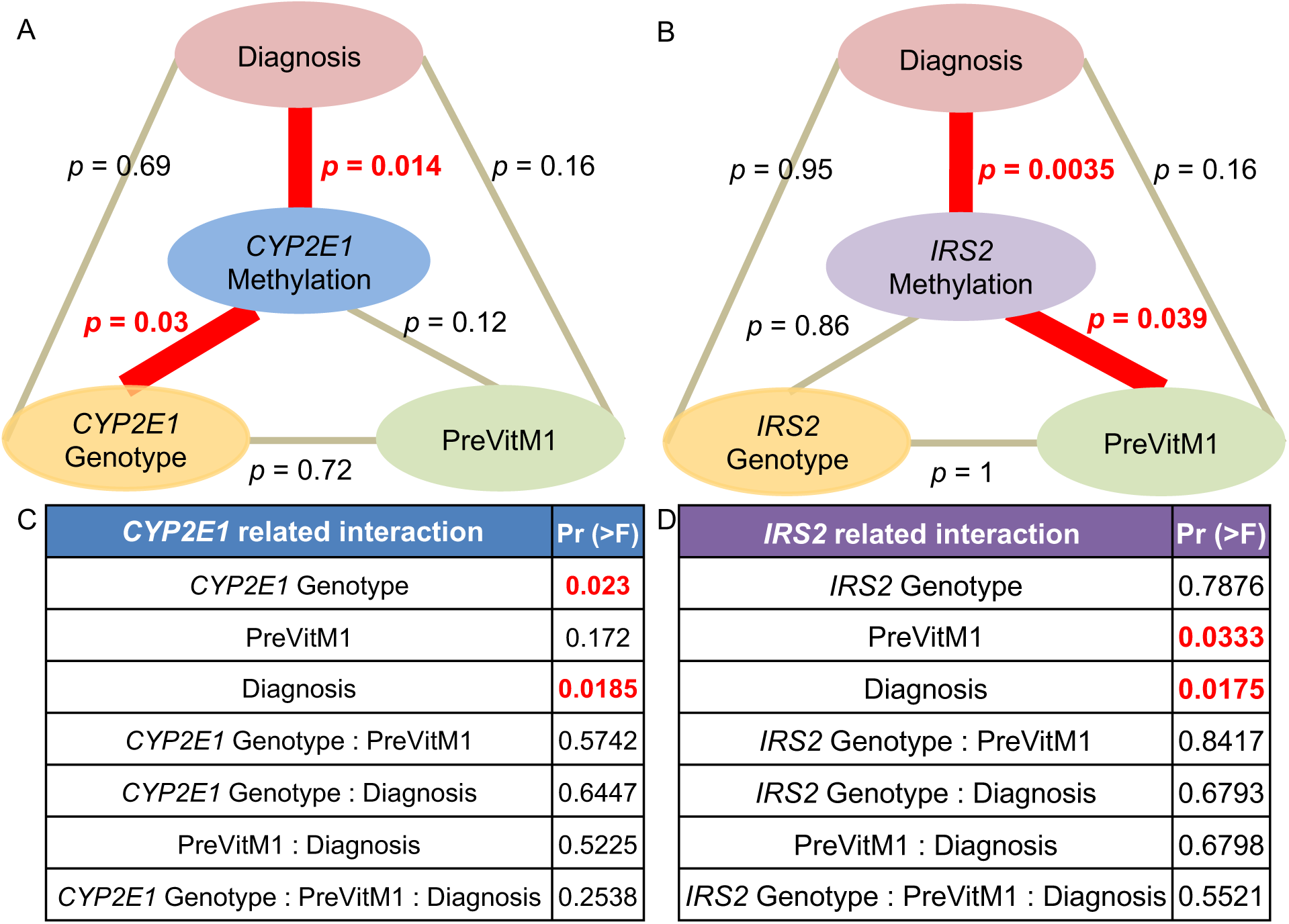
*CYP2E1* and *IRS2* DMR associations and interactions between diagnosis, genotype, and preconception prenatal vitamin use. For **A** and **B**, diagnosis, genotype and PreVitM1 variables were tested for association with methylation separately by two-tailed t-test (or ANOVA for *CYP2E1* genotype) with *p*-value listed at each line. Between each factor, Pearson’s Chi-Squared tests were performed with the *p*-value listed at each line. Significant associations were shown with a bold red line. For **C** and **D**, two-way interactions and three-way interaction were considered by using an ANOVA model to test association among three factors, diagnosis, genotype, and PreVitM1 to methylation at two genome-wide significant DMR. **A**. *CYP2E1* DMR methylation was significantly associated with *CYP2E1* genotype (rs1536828) and diagnosis. **B**. *IRS2* DMR methylation was significantly associated with diagnosis and PreVitM1. **C**. There was a significant association on *CYP2E1* genotype (rs1536828), and diagnosis with *CYP2E1* DMR methylation after considering interaction terms. **D**. Both diagnosis and PreVitM1 were significantly associated with *IRS2* DMR methylation with interaction terms considered.

## Discussion

This study utilized whole methylome analysis of prospectively stored placenta samples in a high risk ASD cohort to bioinformatically identify novel gene loci that were able to discriminate child outcome at age three. This unbiased analysis of ASD differentially methylated regions in placenta tissue resulted in several novel findings. First, the 596 genes identified from 400 placental ASD DMRs significantly overlapped with genetic risk for ASD from curated databases and gene functions in neurons. Second, two genome-wide significant placental ASD DMRs at *CYP2E1* and *IRS2* were discovered that were validated by pyrosequencing and also overlapped with ASD-associated genetic variation and gene expression changes. Lastly, we investigated genotype and nutrient factors correlating with methylation at *CYP2E1* and *IRS2*, demonstrating specific effects for *cis* genotype and diagnosis at *CYP2E1* and prenatal vitamin use at *IRS2*. These results therefore suggest that DNA methylation patterns in placenta provide a direct link between genetics, environment, and fetal epigenetic programming, which can reflect early development relevant to the complex etiology of ASD. The epigenomic signature of ASD in placenta also provides important insights into gene functions, pathways, gene-environment interactions, and potential biomarkers that may be useful in improving early detection of ASD.

This study is the first to identify 400 potential ASD DMRs that distinguish between ASD and TD placenta samples and highlights specific locations and gene functions of differentially methylation in placental samples from children with ASD. First, these placental ASD DMRs were highly enriched around transcription start sites and H3K4me3 marks that are clear marks of gene regulatory functions (Carninci et al., 2006; Koudritsky & Domany, 2008; Portales-Casamar et al., 2007; Yang, Bolotin, Jiang, Sladek, & Martinez, 2007). Furthermore, gene ontology analysis of the 596 genes mapped to placental ASD DMRs pointed to enriched gene functions in transcription, neuron fate, and embryonic development, which were expected based on previous studies (Dapretto et al., 2006; Geschwind & Levitt, 2007; Schroeder et al., 2016). Genes with ASD DMRs in both placenta and brain were enriched for Wnt and cadherin signaling pathways. Wnt signaling is important in embryogenesis, tissue regeneration, and neurodevelopment (Katoh & Katoh, 2006; Logan & Nusse, 2004; Nusse & Clevers, 2017), while cadherin signaling plays a vital role in connecting major intracellular signaling pathways with adhesion protein complexes (Klezovitch & Vasioukhin, 2015; Yap & Kovacs, 2003). Our results therefore complement previous studies that have shown the importance of Wnt and cadherin pathways in the etiology of ASD (Betancur, Sakurai, & Buxbaum, 2009; Kalkman, 2012; Krey & Dolmetsch, 2007). We also replicated our previous finding of differential methylation at *DLL1* in ASD placentas (Schroeder et al., 2016) (**Supplementary Table 2**). *DLL1* encodes a ligand of Notch, activated by Wnt signaling (Hofmann et al., 2004).

When overlapped with datasets of genetic risk for neurodevelopmental disorders including ASD and intellectual disability (Abrahams et al., 2013; Iossifov et al., 2014; Sanders et al., 2015), placenta ASD DMRs were significantly enriched for ASD but not for intellectual disability genetic risk, illustrating the specificity of the ASD DMRs identified in our study. The highest overlap of ASD DMR associated genes was with the SFARI high confidence genes, including *KMT2A, MYT1L*, and *TBR1* (Abrahams et al., 2013). *KMT2A* is expressed in brain and placenta and encodes for a transcriptional coactivator (lysine methyltransferase 2A) that modulates H3K4 methyltransferase activity, specifically the transfer of methyl groups from S-adenosylmethionine to lysine residues on histones (Allis et al., 2007; Shilatifard, 2008) and has been previously implicated in brain development (E. Shen, Shulha, Weng, & Akbarian, 2014; Vallianatos & Iwase, 2015). *MYT1L* encodes for zinc finger transcription factor that functions in the developing mammalian central nervous system and is associated with neurodevelopmental disorders (Blanchet et al., 2017; Wang et al., 2010). *TBR1* (T-box, brain, 1) is a transcription factor which is vital for vertebrate embryo development, neuron migration and differentiation (Bedogni et al., 2010; Englund et al., 2005). In addition, our two genome-wide significant ASD DMR associated genes, *CYP2E1* and *IRS2*, were both on the list of “high confidence” genetic risk for ASD (Sanders et al., 2015).

Four genes were found to be in common between placenta ASD DMRs, brain ASD DMRs and SFARI genetic risk, specifically *GADD45B, MC4R, PCDH9* and *TBL1XR1* (Betancur et al., 2009; Garbett et al., 2008; Orlik & Halawa, 2016; Tabet et al., 2014). *GADD45B* (growth arrest and DNA-damage-inducible, beta) responds to environmental stress through JNK pathway induced DNA demethylation of neurogenesis and synaptic plasticity at gene promoters (Garbett et al., 2008; Ma et al., 2009; Sultan, Wang, Tront, Liebermann, & Sweatt, 2012). *MC4R* encodes the membrane-bound melanocortin 4 receptor, implicated in hormone and cell growth pathways in obesity and insulin resistance (Chambers et al., 2008; Orlik & Halawa, 2016; Yeo et al., 1998). *PCDH9* (protocadherin 9) is a cadherin signaling pathway gene with specific signaling function at neuronal synaptic junctions (Betancur et al., 2009; Bruining et al., 2015). *TBL1XR1* encodes the nuclear receptor corepressor transducing beta like 1 X-linked receptor 1 that binds to histone deacetylase 3 (HDAC 3) complexes in neuron development (Gonzalez-Aguilar et al., 2012; Pons et al., 2015; Tabet et al., 2014). A GWAS noise reduction (GWAS-NR) method to correct for false-positive association with ASD identified cadherin and signaling transduction pathways that included *PCDH9* and *IRS2* as high confidence ASD genes (Hussman et al., 2011).

Our study identified novel methylation differences at *CYP2E1* and *IRS2*, which exhibited genome-wide significant differences between ASD and TD. Both *CYP2E1* and *IRS2* are identified as ASD genetic risk genes in multiple databases related to ASD genetic risk across different tissues and populations (Vogel Ciernia et al., 2018; Sanders et al., 2015). Both *CYP2E1* and *IRS2* DMRs are located close to the TSS site at CpG shore intragenic regions, which is also consistent with the enrichment for TSS flanking regions and H3K4me3 promoter marks in the 400 ASD DMRs. These results are consistent with the developmental dynamics of H3K4me3 marks in human prefrontal cortex, which have been observed to be altered in ASD (Cheung et al., 2010; Shulha et al., 2012). Structural variants and SNPs in cis-regulatory elements also showed significant contribution to ASD (Brandler et al., 2018; W. Sun et al., 2016).

*CYP2E1* encodes a member of the cytochrome P450 superfamily that is involved in the metabolism of drugs, including analgesics like acetaminophen, fatty acids such as arachidonic acid, and a range of chemical toxins, including halogenated hydrocarbons, benzene, and its activity is inducible by drugs, alcohol, and xenobiotics. It thus has an important role in drug bioavailablity (Gonzalez, 1988; Koop, 1992; Rasheed, Hines, & McCarver-May, 1996; Traglia et al., 2017). Previous studies showed those proteins essential for embryonic development in human, rat and zebrafish (Jukka Hakkola et al., 1996; S. M. Jones, Boobis, Moore, & Stanier, 1992; Kishida & Callard, 2001; Ko, Choi, Green, Simmen, & Simmen, 1994; Majdic, Sharpe, O’Shaughnessy, & Saunders, 1996). We observed a significant association between methylation and *cis* genotype at the *CYP2E1* DMR, a finding which is consistent with the identification of this locus in a screen for human metastable epialleles variability between individuals (Silver et al., 2015). In addition, in immune models of ASD, maternal interleukin-6 (IL6) crosses the placenta, disrupting development of hippocampal spatial learning (Boksa, 2010; Jonakait, 2007; Krakowiak et al., 2012). Previous studies showed that IL6 inhibits *CYP1A1, CYP1A2* and *CYP2E1* expression (Abdel-Razzak et al., 1993; J. Hakkola, Hu, & Ingelman-Sundberg, 2002; Jover, Bort, Gómez-Lechón, & Castell, 2002; Patel et al., 2014), consistent with the lower methylation and expression levels in ASD versus TD observed in our study. In addition, *CYP2E1* expression is transcriptionally regulated by the JAK2/STAT3 pathway, providing a potential convergent pathway with *IRS2* (**Fig. 9**) (Patel et al., 2014).

**Figure 9.**
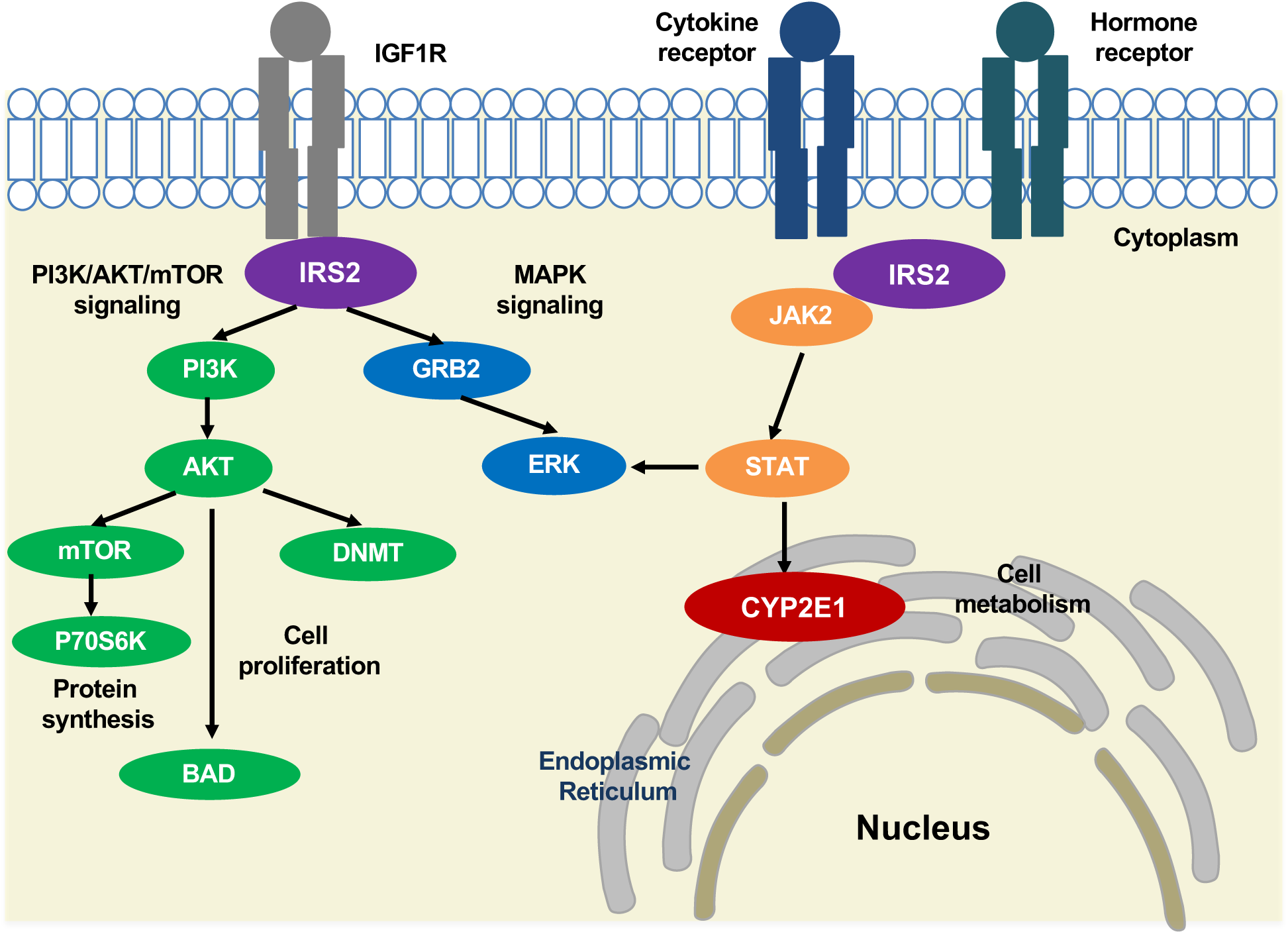
Potential pathway convergence of proteins encoded by both ASD DMRs. IRS2 interacts with transmembrane protein insulin-like growth factor receptor (IGF1R) at the intracellular membrane, resulting in activation of the PI3K/AKT/mTOR and MAPK signaling pathways (Machado-Neto et al., 2018) involved in protein synthesis, cell proliferation and gene expression (Archuleta et al., 2009; Machado-Neto et al., 2018; Patti et al., 1995; Tamburini et al., 2008; Velloso et al., 1996). An AKT-mediated ubiquitin pathway leads to *de novo* DNA methylation changes by DNMT (FANG et al., 2015; Lin & Wang, 2014). IRS2 also interacts with cytokine and hormone receptors and induces JAK2/STAT3 signaling (Carvalheira et al., 2003; Machado-Neto et al., 2018; Saad et al., 1996, 1995). STAT activation leads to *CYP2E1* localization at the endoplasmic reticulum, changing cellular metabolism (Patel et al., 2014).

*IRS2* encodes for insulin receptor substrate 2, a cytoplasmic signaling molecule that mediates the effects of insulin and insulin-like growth factor 1 (*IGF1*) (Park et al., 2016; Withers et al., 1998) and cytokine receptors (Ihle, 1995; Rui, Yuan, Frantz, Shoelson, & White, 2002; X. J. Sun et al., 1995). IRS2 has a phosphotyrosine-binding domain which contributes to the intracellular affinity to cell membrane receptors (**Fig. 9**) (Eck, Dhe-Paganon, Trüb, Nolte, & Shoelson, 1996; Machado-Neto, de Melo Campos, & Traina, 2018; Schlessinger & Lemmon, 2003; White, 1998). PI3K/AKT/mTOR and MAPK signaling pathways are linked with IRS2 in the regulation of protein synthesis and cell proliferation (Archuleta et al., 2009; Machado-Neto et al., 2018; Patti et al., 1995; Tamburini et al., 2008; Velloso et al., 1996). When activated by cytokine and hormone receptors, IRS2 stimulates JAK2, leading to STAT and MAPK signaling activation (Carvalheira, Ribeiro, Folli, Velloso, & Saad, 2003; Machado-Neto et al., 2018; Saad, Carvalho, Thirone, & Velloso, 1996; Saad, Velloso, & Carvalho, 1995; Velloso et al., 1996). The insulin-like growth factors (IGF1) pathway, which includes IRS2, also mediates *de novo* DNA methylation by DNA methyltransferase (DNMT) through AKT (Fang et al., 2015; Lin & Wang, 2014). This pathway may explain why methylation at the *IRS2* DMR was sensitive to maternal prenatal vitamin intake, since *IRS2* stimulates the mTOR (mechanistic target of rapamycin) pathway, which responds to nutrients and growth factors signaling to regulate protein synthesis (Gulati & Thomas, 2007; M. Laplante & Sabatini, 2013; Mathieu Laplante & Sabatini, 2012; Shimobayashi & Hall, 2014). Previous studies have shown that maternal folate alters amino acid transport activity in the placenta, resulting in affecting fetal growth by the mTOR signaling pathway (Huang & Fingar, 2014; Shimobayashi & Hall, 2014). Our analysis shown that placental differentially methylated gene loci associated with prenatal vitamin intake were also highly enriched for functions in fetal growth and development. We showed that high *IRS2* methylation is significantly associated with ASD and mothers who did not take prenatal vitamins before conception, suggesting that sufficient folate levels around placental implantation may be protective for ASD through IRS2-mTOR signal transduction. The link between epigenetic alterations in *IRS2* and risk for ASD is particularly intriguing given a growing body of epidemiologic evidence demonstrating higher ASD risk in offspring born to mothers who experienced diabetes during pregnancy, including some very large and methodologically sound studies with clinical diagnoses of both maternal diabetes and child ASD (Li et al., 2016; Xiang et al., 2018; G. Xu, Jing, Bowers, Liu, & Bao, 2014).

We did not observe any significant associations between other potential cofounders such as maternal age, pregnancy BMI, or gestational age at birth and ASD diagnosis in the MARBLES study (Schmidt et al., 2016). Cell type heterogeneity in the placenta may complicate the interpretation of our results, however, our previous study did not detect differences in methylation levels by placental region at specific gene loci (Schroeder et al., 2016). Other potential limitations of our study include the relatively small sample size and sequencing depth. This study serves as a proof-of-principle that placenta methylation patterns detected by WGBS may be informative in ASD. Replication with additional samples, other similar prospective cohorts, and improved sequencing and bioinformatic strategies will be important in future studies.

In conclusion, we identified two high confidence genes differentially methylated in ASD from an unbiased analysis of DNA methylation in placenta from high-risk pregnancies and investigated possible genetic and environmental modifiers of methylation at both loci. Methylation levels at the *CYP2E1* DMR were associated with genotype, while the methylation levels at the *IRS2* DMR were associated with prenatal vitamin use. Our results are consistent with a previous study using the Illumina 450K array, which showed that both genetic and environmental effects influence DNA methylation levels (Hannon et al., 2018). Placenta reflects the essential interface between the fetus and mother, mediating the impacts of endocrine and growth factors in the maternal environment on fetal development (Koukoura, Sifakis, & Spandidos, 2012; Zeltser & Leibel, 2011). Both *CYP2E1* and *IRS2* are related to protein synthesis, cell proliferation, and cell metabolism, consistent with previous studies of convergent gene pathways in ASD (Vogel Ciernia et al., 2018; Sanders et al., 2015; Voineagu et al., 2011; Xu et al., 2012; Zhang et al., n.d.). These results therefore provide evidence that placental methylation levels reflect the intersection of genetic and environmental risk and protective factors that are expected to be useful for early intervention and prevention of ASD.

## Materials and Methods

### MARBLES study design, sample selection, and DNA isolation

The Markers of Autism Risk in Babies: Learning Early Signs (MARBLES) study design was described in a previous publication (Hertz-Picciotto et al., 2018). In MARBLES, mothers of at least one child with confirmed ASD who were pregnant or planning a pregnancy were recruited in the Northern California area. Inclusion criteria for the study were: 1) mother or father has one or more biological child(ren) with ASD; 2) mother is 18 years or older; 3) mother is pregnant; 4) mother speaks, reads, and understands English sufficiently to complete the protocol and the younger sibling will be taught to speak English; 5) mother lives within 2.5 hours of the Davis/Sacramento, California region at time of enrollment. With shared genetics, the next child has a 15-fold higher risk for developing ASD compared to the general population (Hertz-Picciotto et al., 2018). Demographic, diet, lifestyle, environmental, and medical information were prospectively collected through telephone-assisted interviews and mailed questionnaires throughout pregnancy and the postnatal period. Infants received standardized neurodevelopmental assessments beginning at 6 months and concluding at 3 years of age (Hertz-Picciotto et al., 2018). Diagnostic assessments at 3 years included the gold standard Autism Diagnostic Observation Schedule (ADOS) (Lord, Risi, Lambrecht, Cook Jr, et al., 2000; Lord, Risi, Lambrecht, Cook, et al., 2000), the Autism Diagnostic Interview-Revised (ADI-R) (Lord, Rutter, & Le Couteur, 1994), and the Mullen Scales of Early Learning (MSEL) (Mullen, 1995). Participants were classified into outcome groups including ASD and Typical Development (TD), based on a previously published algorithm that uses ADOS and MSEL scores (Chawarska et al., 2014; Ozonoff et al., 2014). Children with ASD outcomes have scores over the ADOS cutoff and meet DSM-5 criteria for ASD. Children with TD outcomes have all MSEL scores within 2.0 SD and no more than one MSEL subscale that is 1.5 SD below the normative mean and scores on the ADOS at least three more points below the ASD cutoff. This study utilized 41 male MARBLES placenta samples, including 20 samples from children later diagnosed with ASD and 21 children determined to have TD, matched for enrollment time frame and date of birth. DNA was isolated from 50–100 mg frozen placental tissues (20 ASD and 21 TD) using the Gentra Puregene tissue kit (Qiagen).

### Whole Genome Bisulfite Sequencing (WGBS)

Raw sequencing data (fastq files) were published previously (Schroeder et al., 2016). Briefly, WGBS libraries were made with the sonicated genomic DNA (around 300 bp) and ligated with methylated Illumina adapters using NEB’s NEBNext DNA library prep kit (Schroeder et al., 2013, 2016). The library was bisulfite converted using EZ DNA Methylation lighting kit (Zymo), amplified for 12 cycles using PfuTurbo Cx Hotstart DNA Polymerase (Agilent) and purified with Agencourt AMPure XP Beads (Beckman Coulter). The quality and quantity of libraries were measured on a Bioanalyzer (Agilent) and sequenced on Illumina HiSeq 2000 with each sample per single lane. Reads after trimming were uniquely mapped to human reference genome (hg38) as described previously using BS-Seeker2 on average 1.6X genome converge with 99.3% bisulfite conversion efficiency (measured through the percentage of non-CpG cytosines that were unconverted) (Dunaway et al., 2016; Guo et al., 2013; Schroeder et al., 2013).

### ASD Differentially Methylated Regions (DMRs) and genome-wide significant DMRs

DMRs were called as described in previous publications (Coulson et al., 2018; Dunaway et al., 2016; Mordaunt, Shibata, et al., 2018) using the default settings. In this case, each ASD DMR contained greater than 10% methylation difference between ASD and TD samples at least three CpGs within 300 base pairs (bp) and a *p*-value < 0.05. Background regions were defined using the same conditions as DMRs but without any percent methylation filters to identify all possible DMR locations based on CpG density and sample sequencing coverage. Hypermethylated ASD DMRs were defined as higher percent methylation in ASD versus TD, while hypomethylated ASD DMRs was were defined as lower percent methylation in ASD versus TD samples. Genome-wide, significant DMRs were identified based on a family-wide error rate (FWER) < 0.05, determined by permuting the samples 1000 times by chromosome, and counting the number of null permutations with equal or better DMRs ranked by number of CpGs and areaStat (Box, 1980).

### Hierarchical clustering and principal component analysis (PCA)

Methylation was extracted at each ASD DMR for every sample. Percent methylation of each sample was normalized to the mean methylation of each ASD DMR. ASD DMRs were grouped by Ward’s Method of hierarchical clustering (Wilks, 2011). Principal component analysis was performed on methylation at all ASD DMRs across all samples using the prcomp function and ggbiplot package in R. The ellipses for each group were illustrated as the 95% confidence interval. The lack of overlapping ellipses for ASD and TD samples indicated significant methylation difference in ASD DMRs between groups (*p* < 0.05).

### Assignment of DMRs to genes and relative location to TSS

Genes were assigned to DMRs using the Genomics Regions Enrichment of Annotations Tool (GREAT) on the default association settings (5.0 kilo-base (kb) upstream and 1.0 kb downstream, up to 1000.0 kb max extension) (McLean et al., 2010). The distance (kb) was calculated from the ASD DMRs, hypermethylated ASD DMRs, hypomethylated ASD DMRs and background regions to transcription start site (TSS) of the GREAT assigned gene. The gene length was calculated for both placental ASD DMR genes and all genes in human genome and tested for potential distribution differences by Pearson’s chi-squared test.

### Gene Ontology Term and Pathway Enrichment Analysis

Gene ontology (GO) analysis was done using PANTHER (Protein Analysis Through Evolutionary Relationships) overrepresentation test, with the GO Ontology database (Ashburner et al., 2000; The Gene Ontology Consortium, 2017) and Fisher’s exact test with false discovery rate (FDR) multiple test correction. GO term enrichments were presented by the hierarchical terms rather than specific subclass functional classes, as described previously (Mi, Muruganujan, & Thomas, 2012; Thomas et al., 2003).

### Tests for ASD DMR Enrichments

All tests of enrichment for ASD DMRs were compared to a set of all possible background regions that are calculated in the DMR analysis pipeline. Enrichment tests for placenta ASD DMRs associated genes and published gene lists were done using the GeneOverlap R package which implements Fisher’s exact test and adjusted for FDR correction (L. Shen et al., 2013). \**p* < 0.05, \*\**p* < 0.01, \*\*\**p* < 0.001 by Fisher’s exact test with FDR corrected. Brain cortex (BA9) ASD DMRs were defined as either a 5% or 10% methylation difference between ASD and TD and were described previously using the same method as placenta ASD DMRs (Vogel Ciernia et al., 2018). The SFARI (Simons Foundation Autism Research Initiative) database was used for the five categories of ASD risk genes (https:gene.sfari.org/database/gene-scoring/) (Abrahams et al., 2013). High effect ASD risk gene lists were also identified from Sanders *et al*. (Sanders et al., 2015). Likely gene-disrupting (LGD) recurrent ASD mutations and missense mutation on *de novo* mutations were obtained from Iossifov *et al*. (Iossifov et al., 2014). Gene lists on intellectual disability (ID) were obtained from Gilissen *et al*. (Gilissen et al., 2014). Alzheimer’s disease GWAS gene lists were extracted from SNPs showing association with Alzheimer’s disease (*P* ≤ 1×10^−3^) (Harold et al., 2009). Lung cancer GWAS gene lists were acquired from Landi *et al*. (Landi et al., 2009). The random genes category contains the same number of regions as the placenta ASD DMRs to serve as a specificity control. ASD DMRs were examined for enrichment with known chromatin marks compared to the background using LOLA R package with two-tailed Fisher’s exact test after FDR correction (Sheffield & Bock, 2016). Placenta histone marks H3K4me1, H3K4me3, H4K9me3, H3K36me3, H3K27me3 and H3K27ac were extracted from ENCODE (Encyclopedia of DNA Elements) placenta ChIP-seq dataset (ENCODE Project Consortium, 2012; Sloan et al., 2016). ASD DMRs were also analyzed for overlap with chromatin states predicted by chromHMM, which use histone modification ChIP-seq data to separate the genome into 15 functional states in the Roadmap Epigenomics Project using a Hidden Markov Model (Ernst & Kellis, 2017; Kundaje et al., 2015). For promoters, chromHMM separates active transcription start site (TssA), TSS flank (TssAFlnk), bivalent TSS (TssBiv), and bivalent TSS flank (BivFlnk) states. For enhancers, genic enhancer (EnhG), enhancer (Enh), and bivalent enhancer (EnhBiv) are the different states. Human CpG island locations were extracted from UCSC genome browser (Kent et al., 2002). CpG island shores were defined as 2 kb flanking regions on both sides of CpG island. CpG island shelf was characterized as 2 kb flanking regions on both sides of CpG island shore, not including CpG island or CpG island shore. CpG island “open sea” includes all genomic regions except CpG island, CpG island shore and CpG island shelf. A custom R script was used to generate the locations of CpG islands (https:github.com/Yihui-Zhu/AutismPlacentaEpigenome).

### Pyrosequencing

Genomic DNA (500 ng) was bisulfite converted using the EZ DNA Methylation kit (Zymo). Amplification and sequencing primers were designed using the PyroMark Assay Design Software 2.0 (Qiagen). DMRs were amplified using the PyroMark PCR kit (Qiagen). Pyrosequencing of 13 CpG sites at *CYP2E1* gene, and 11 CpG sites in human *IRS2* gene was performed in triplicate. Pyrosequencing was performed on a Pyromark Q24 Pyrosequencer (Qiagen) with the manufacturers recommended protocol. Enzyme, substrate, and dNTPs were from the Pyromark Gold Q24 Reagents (Qiagen) and the methylation levels were analyzed using Pyromark Q24 software.

*CYP2E1* related DMR pyrosequencing primers:

Forward: GGTGTTTTGTTTTGGGGTTGA

Reverse: ACCCATTCAATATTCACAACAATC (5’ Biotin)

Sequencing: GGTTGATGATGGGGA

Amplification region: chr10: 133527817 – 133527938 (hg38)

*IRS2* related DMR pyrosequencing primers:

Forward: TTAGGAATATAGGGAAAGGTGAAAGT

Reverse: CCACCCATTCACCCATTCTA (5’ Biotin)

Sequencing: GGGAAAGGTGAAAGTT

Amplification region: chr13: 109781623 – 109781794 (hg38)

### Gene Expression in Umbilical Cord Blood

Data for gene expression assessed by Affymetrix Human Gene 2.0 array were extracted a previous publication on umbilical cord blood from subjects in the MARBLES study (GEO ID: GSE123302) (Mordaunt, Park, et al., 2018). Placenta and cord blood were collected at the same time period in the same study. Raw intensity values from cord blood samples were normalized by RMA and data from 70 male samples were extracted, including 30 ASD and 40 TD samples. Normalized expression was examined at the only probe annotated to *CYP2E1* (16711001) and the only probe annotated to *IRS2* (16780917). Analysis was done on those two probes with 70 samples on the normalized matrix data.

### Western Blot

In Western blot experiments, placental proteins were isolated with RIPA buffer containing 10mM Tris-Cl (pH 8.0), 1mM EDTA, 1% Triton X-100, 0.1% sodium deozydholate, 0.1% SDS, 140mM NaCl, 1mM PMSF and complete protease inhibitors (ThermoFisher), incubated at 37°C for 30 minutes, sonicated and heated at 95°C for 5 min. BCA (Bicinchoninic Acid) protein assay (ThermoFisher) was used to determine protein concentration. Protein samples (20–30 ug) were resolved on 4–20% tris-gylcine polyacrylamide gels (Biorad). Proteins were separated and transferred to nitrocellulose membranes for 60 minutes at a constant voltage of 100. The membranes were blocked in Odyssey Blocking Buffer (PBS) (Licor, 927–40000) for 40 min. Anti-IRS2 (1:5,000, Cell Signaling, 3089S) and anti-GAPDH (1:10,000, Advanced Immunochemical, Inc., 2-RGM2) were incubated with the membrane with Odyssey Blocking Buffer containing 0.2% Tween overnight at 4°C. Membranes were washed with 1 X PBS (Phosphate-buffered saline) containing 0.2% Tween and then incubated with secondary antibodies, IRDye 800CW Donkey anti-Mouse IgG (1:50,000, Licor, 926–32212) and IRDye 680RD Donkey anti-Rabbit IgG (1:50,000, Licor, 926–68073) for 1 hour. Membranes were scanned using a Licor Odyssey infrared imaging system based on the manufacturer’s guidance (with resolution: 84; quality: medium, 600-channel: 6; 800-channel: 5). Relative protein quantification was done using the ImageJ software program (Rueden et al., 2017; Schneider, Rasband, & Eliceiri, 2012) in densitometry mode. IRS2 signals were normalized to GAPDH (Glyceraldehyde 3-phosphate dehydrogenase) for each sample.

### Sanger Sequencing

PCR amplification was performed on each sample using PCR 10x buffer, 25 mM MgCl_2_, 5 M betaine, 10 mM dNTPs, DMSO, and HotStart Taq (Qiagen). Each PCR program was unique to the region being amplified with specific primers. The PCR product was then resolved by gel electrophoresis using a 1% Agarose gel in 1 X TE to later be extracted using the gel extraction kit (Qiagen) based on the default protocol. After DNA quantitation by NanoDrop, the samples were sent to the UC Davis Sequencing Facility for sequencing on the 3730 Genetic Analyzer (Applied Biosystems Prism) with DNA sequencing Analysis software v.5.2 (Applied Biosystems Prism). The sequencing results were assembled and analyzed using CodonCode Aligner version 7.0 (CodonCode).

*CYP2E1* related SNP (rs943975, rs1536828) primers:

Forward: CTACAAGGCGGTGAAGGAAG

Reverse: CCCATCCCCATAAACTCTCC

*IRS2* related SNP (rs943975) primers:

Forward: TTAGGAATATAGGGAAAGGTGAAAGT

Reverse: CCACCCATTCACCCATTCTA

### Maternal Prenatal Vitamin Use and Timing

Maternal prenatal vitamin information and timing of maternal intake for 6 months before and each month during the pregnancy were recorded though telephone interviews and/or questionnaires as previously describes (Hertz-Picciotto et al., 2018). Mothers who took prenatal vitamins in the first month pregnancy or not were grouped into PreVitM1 Yes/No. Mothers who took prenatal vitamins beginning from 6 months to 2 months before pregnancy were grouped as “Before Pregnancy”. Mothers beginning prenatal vitamins one month before pregnancy through the second month of pregnancy were grouped as “Near Conception”. Mothers beginning prenatal vitamins from 3 months to 9 months of pregnancy were grouped as “During Pregnancy”.

### Code availability

Custom scripts for WGBS analysis are available at https:github.com/kwdunaway/WGBS_Tools with the instructions. Custom Scripts for DMR finder are available at https:github.com/cemordaunt/DMRfinder with the instructions. The rest of code and scripts for each figure and tables are available at https:github.com/Yihui-Zhu/AutismPlacentaEpigenome.

### Data availability

WGBS data were previously published, Gene Expression Omnibus (GEO) accession number GSE67615 (Schroeder et al., 2016). The rest of the relevant data and information are included in supplementary tables.

## Supporting information

Supplementary Figures and List of Supplementary Tables

## Acknowledgements

We would like to thank members of the LaSalle lab and UCD Children’s Center for Environmental Health for helpful discussions, and the MARBLES study participants. This work used the Vincent J. Coates Genomics Sequencing Laboratory at UC Berkeley, supported by NIH S10 Instrumentation Grants S10RR029668 and S10RR027303. This work was supported by R01 ES025574, NIH R01ES021707, NIH P01ES011269 (CCEH), EPA 83543201 (CCEH), DOD AR110194, R01ES020392 (MARBLES), U54HD079125 (IDDRC), P30ES023513 (EHSC) and NIH-UL1-TR000002 (CTSC).

## Competing interests

The authors declared they do not have anything to disclose regarding funding or conflict of interest with respect to this manuscript.

