## Supplementary Figures and List of Supplementary Tables for "Placental DNA methylation levels at *CYP2E1* and *IRS2* are associated with child outcome in a prospective autism study"

**Contents** **Pages**

| Contents | Pages |
|---|---|
| Supplementary Figures | 2 |
| List of Supplementary Tables | 10 |

### Supplementary Figures:

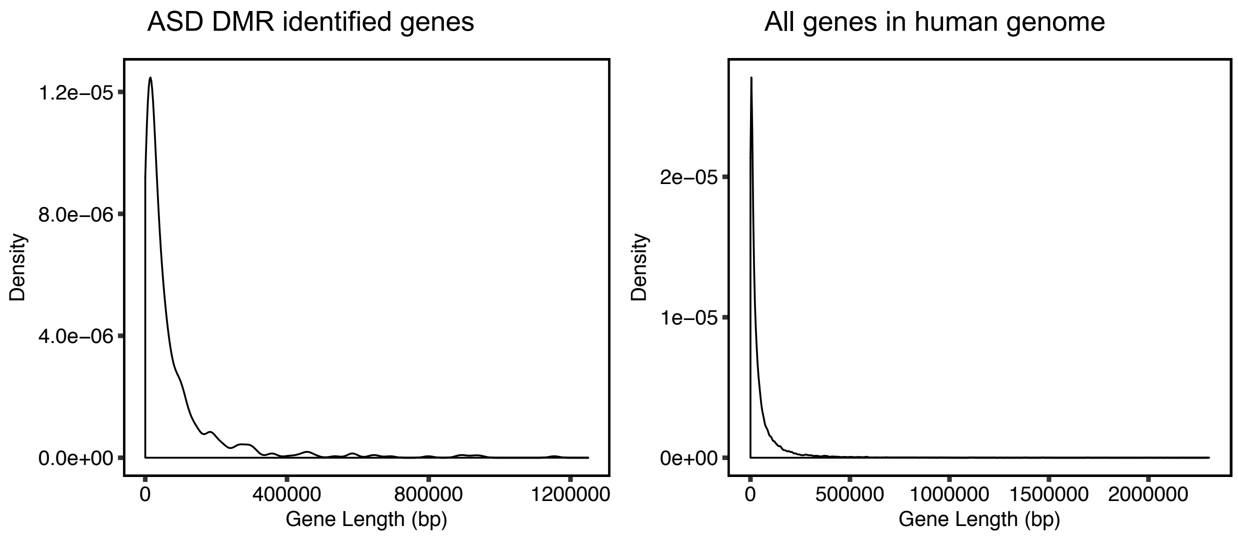


**Supplementary Figure 1: The distribution of gene length for ASD DMR genes was similar to all genes in the human genome.**

The density plot showed the distribution of gene length of ASD DMR identified genes (left) and all genes (right). The x-axis illustrated the gene length in base pairs. The y-axis was the density for gene length. Person's chi-squared test showed no significant difference between two distributions (*p*-value = 0.9994).

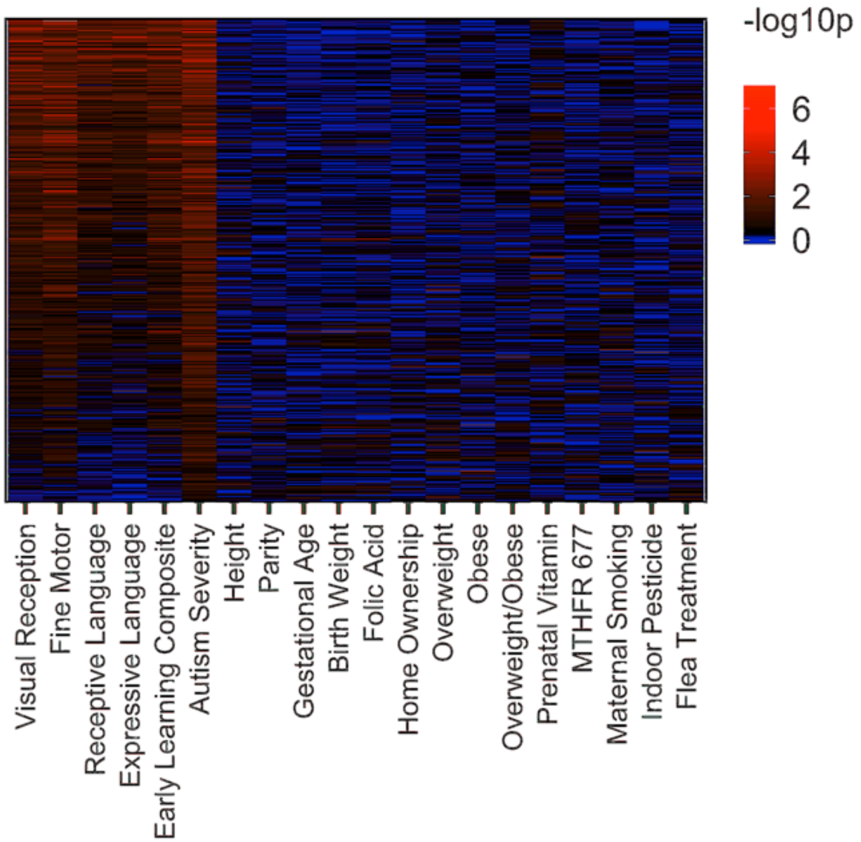


**Supplementary Figure 2: ASD DMRs heatmap by child outcome continuous measurements of cognition and autism severity versus potential cofounding variables.**

The plot shows a heatmap of ASD DMRs (y-axis) and the association of % methylation at each DMR with other measured variables. The first 5 child outcome variables on the x-axis includes 4 sub-categories of Mullen scores as well as composite score and autism severity score from the ADOS. Significant associations are red ($p < 0.05$). While ASD DMRs were highly associated with autism severity and to a lesser degree with early learning Mullen's scores, other potential confounding variables from MARBLES exhibited only rare associations with individual ASD DMRs.

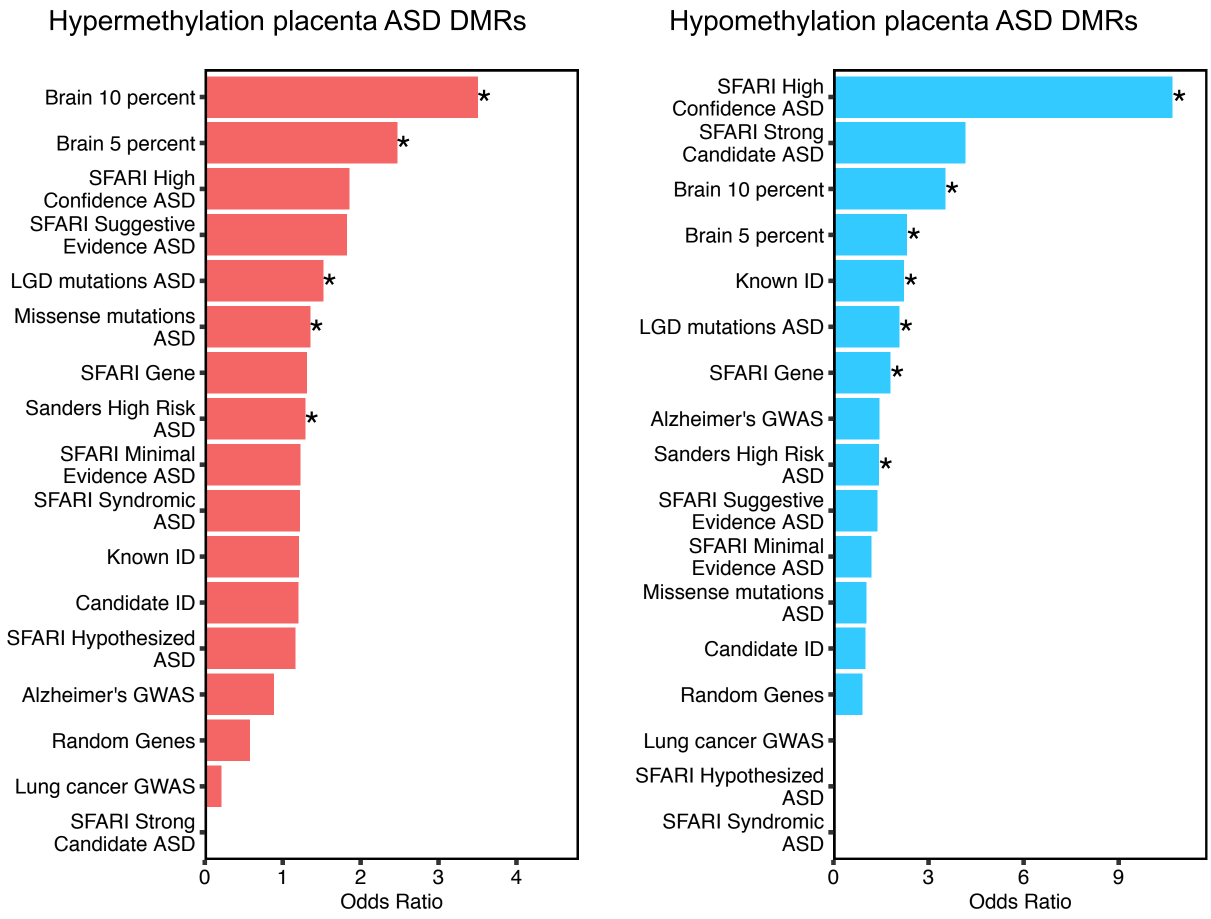


**Supplementary Figure 3: Enrichment test on hyper- versus hypomethylated placenta ASD DMR genes for overlap with ASD risk genes and other databases.** Hypermethylated (left) and Hypomethylated (right) placenta ASD DMRs associated genes overlapped with ASD genetics risk database, other intellectual disability and a random gene list ranked by odds ratio. *$p < 0.05$, **$p < 0.01$, ***$p < 0.001$ by two tailed Fisher's exact test after the FDR correction. SFARI: Simons Foundation Autism Research Initiative (Abrahams et al., 2013), LGD: likely gene disrupting mutation, ASD: autism spectrum disorder, Alzheimer: Alzheimer's Disease, ID: intellectual disability.

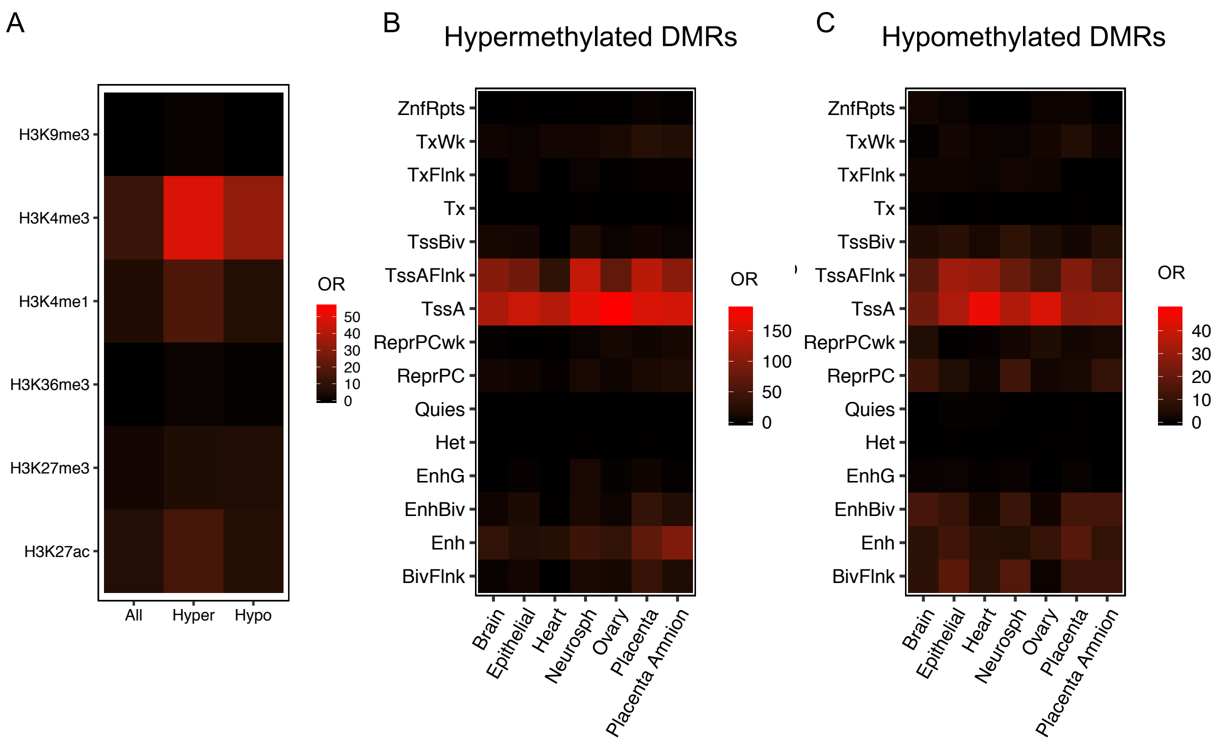


**Supplementary Figure 4: Placenta ASD hyper- and hypomethylated DMRs were both enriched at H3K4me3 regions, active promoters and their flanking regions.**

**A**. Hyper- and hypomethylated placenta ASD DMRs were overlapped with histone modification human placenta ChIP-seq peaks from the Epigenome Roadmap with odds ratio plotted

**B**. Hypermethylated placenta ASD DMRs were tested on chromatin states from the Epigenome Roadmap. The x-axis was different tissue type, the y-axis was represented chromatin states.

**C**. Hypomethylated placenta ASD DMRs were tested on chromatin states from the Epigenome Roadmap.

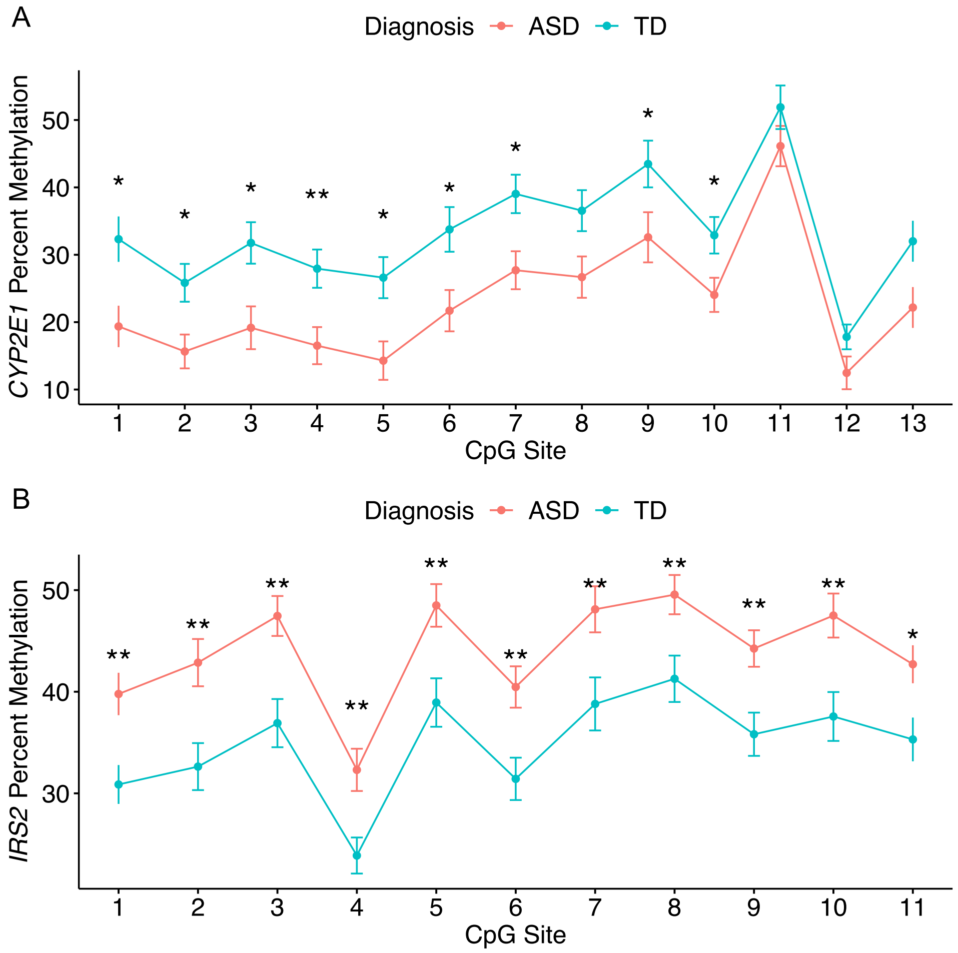


**Supplementary Figure 5: Pyrosequencing results from *CYP2E1* and *IRS2* DMRs for each CpG site.**

The x-axis represents each CpG sites included in pyrosequencing DMR regions. The y-axis plots the percent methylation at each CpG site. The red line showed the average methylation of ASD samples and the blue line represented the average methylations of TD samples and error bars represent the standard error of the mean. Each CpG site was tested on the significance level with FDR corrected for the numbers of CpGs. *$p < 0.05$, **$p < 0.01$. **A**. 13 CpG sites tested at the *CYP2E1* DMRs with 10 of them showed significant association with diagnosis after FDR correction. **B**. All 11 CpG sites at the *IRS2* DMRs showed significant association with diagnosis after FDR correction.

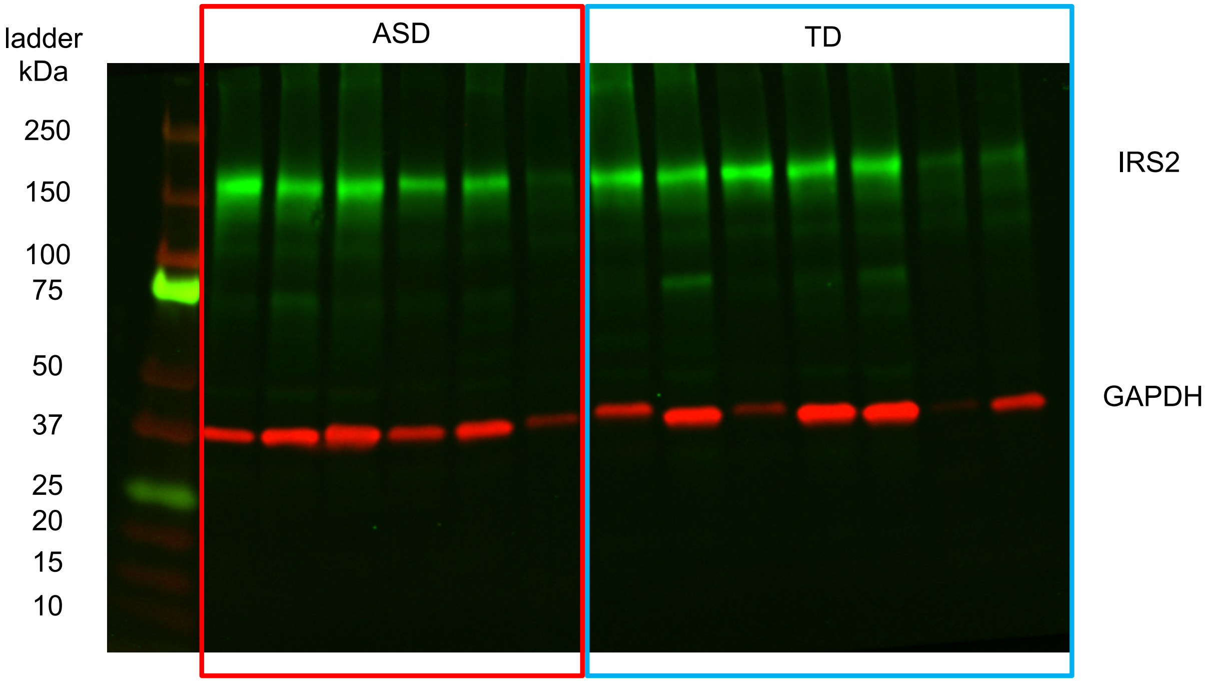


**Supplementary Figure 6: Representative IRS2 Western blot image.**

A representative Western blot image of placental protein extracts detected by anti-IRS2 (green) or anti-GAPDH (red) antibodies. The left column is the protein size ladder.

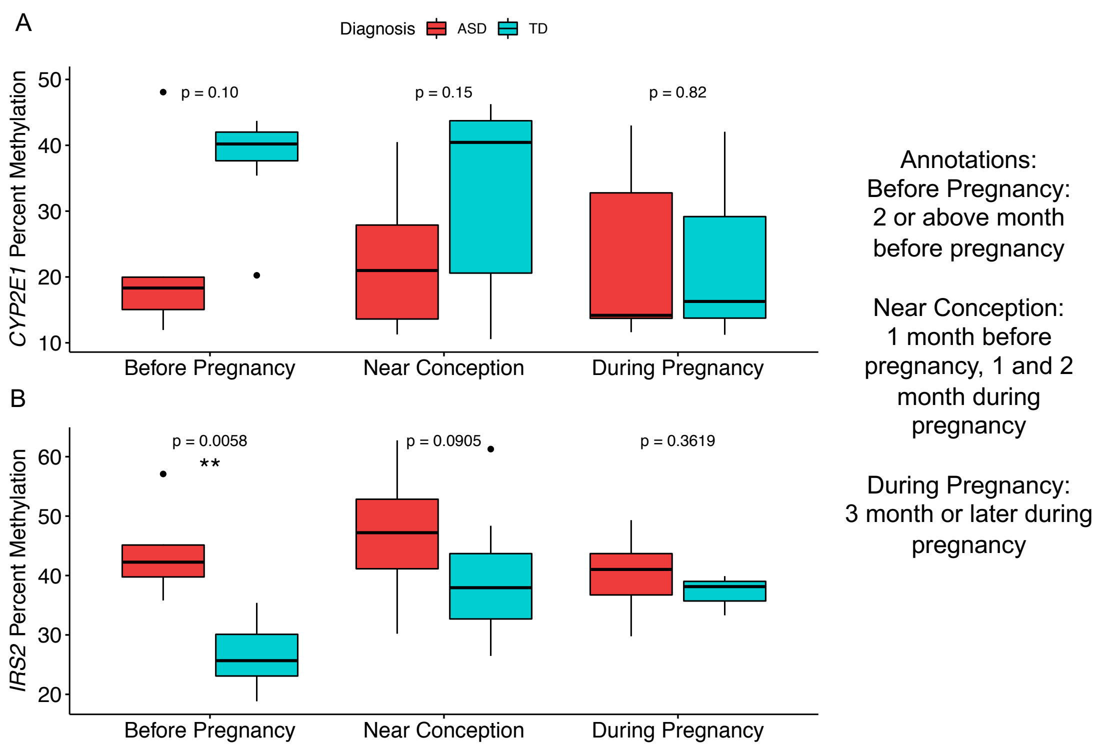


**Supplementary Figure 7: Prenatal vitamins intake prior to pregnancy was protective for placental DNA methylation patterns at *CYP2E1* and *IRS2* DMRs.** Each sample was labelled at the month the mother started taking prenatal vitamins. The time line was separated into three time bins. Mothers who started prenatal vitamin use more than two months before pregnancy were grouped as "before pregnancy". Those who started one month before pregnancy, first, or second month of pregnancy were categorized as "near conception", while "during pregnancy" include those who started taking prenatal vitamins 3 months or later into pregnancy. Two-tailed t-test was done within each category between ASD and TD samples. For both *CYP2E1* (**A**) and *IRS2* (**B**) DMRs, the largest protective effect of prenatal vitamin use on methylation levels was observed in placentas from mothers who started prenatal vitamin use before pregnancy.

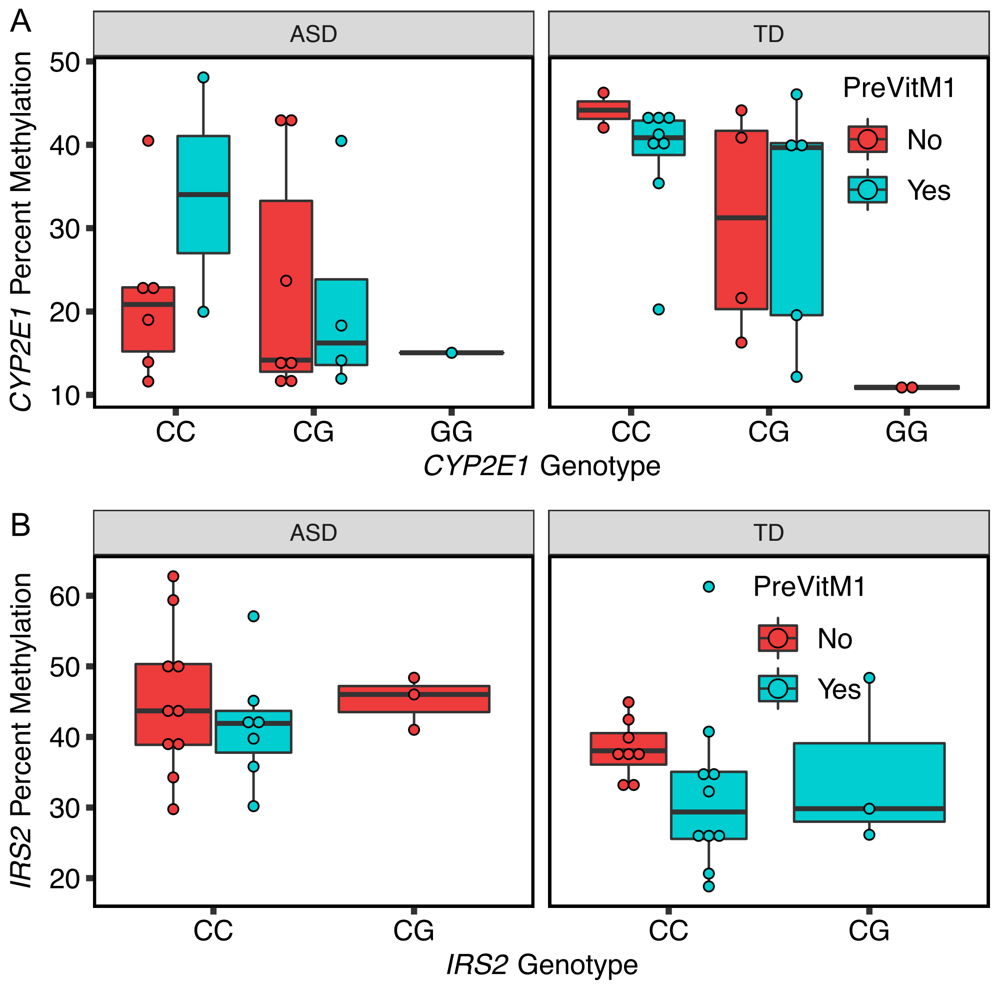


**Supplementary Figure 8: Comparison of methylation levels at *CYP2E1* and *IRS2* DMRs by diagnosis, genotype and preconception prenatal vitamin use.** Pyrosequencing methylation results (y-axis) were separated by diagnosis, ASD and TD and graphed separately by P1 prenatal vitamins use (Yes/No). The x-axis shows the *cis* genotype of a common SNP within each DMR.

**Supplementary Tables:**

**Supplementary Table 1**:

400 differentially methylated regions (DMRs) in placenta that distinguish ASD and TD samples and their association with 597 genes.

**Supplementary Table 2**:

Neurodevelopmental outcomes and additional variables for each placenta sample.

**Supplementary Table 3**:

597 genes associated with ASD DMRs and the distance (bp) to the transcription start site (TSS).

**Supplementary Table 4**:

Gene ontology (GO) analysis of ASD DMR genes by Fisher's exact test after FDR (false discovery rate) correction.

**Supplementary Table 5**:

Overlapping genes between placenta ASD DMR associated genes and other databases, including brain ASD DMR associated genes, ASD genetic risk factors, intellectual disability, Alzheimer's GWAS, and lung cancer GWAS.

**Supplementary Table 6**:

Methylation data from both *CYP2E1* and *IRS2* DMRs for each sample and CpG site (13 CpG sites for *CYP2E1* and 12 CpG sites for *IRS2*). The average represents all CpG sites for each sample. Methods and primers were described in methods sections.

**Supplementary Table 7**:

Sanger sequencing results from *CYP2E1* DMR and *IRS2* DMR on rs943975, rs1536828 and rs9301411.

**Supplementary Table 8**:

376 differentially methylated regions (DMRs) in placenta separated by whether prenatal vitamins were taken or not during the first month of pregnancy. 587 genes were associated with PreVitM1 DMRs. The interaction set of 60 genes with ASD DMRs is also shown.

**Supplementary Table 9**:

Data on whether mothers took prenatal vitamins or not at each month during six months before pregnancy and nine months after conception. “0” stands for “No” and “1” stands for “Yes” in each month.

**Supplementary Table 10**:

Methylation at the *CYP2E1* DMR and *IRS2* DMR was tested for interaction with diagnosis, genotype, and PreVitM1.
